# Multi-omic profiling of primary mouse neutrophils reveals a pattern of sex and age-related functional regulation

**DOI:** 10.1101/2020.07.06.190595

**Authors:** Ryan Lu, Shalina Taylor, Kévin Contrepois, Mathew Ellenberger, Nirmal K. Sampathkumar, Bérénice A. Benayoun

**Affiliations:** Leonard Davis School of Gerontology, University of Southern California, Los Angeles, CA 90089, USA; Graduate program in the Biology of Aging, University of Southern California, Los Angeles, CA 90089, USA; Departments of Pediatrics and of Medicine, Stanford University School of Medicine, Stanford, CA 94305, USA; Department of Genetics, Stanford University, Stanford, CA 94305, USA; UK-Dementia Research Institute, Institute of Psychiatry, Psychology and Neuroscience, Basic and Clinical Neuroscience Institute, Maurice Wohl Clinical Neuroscience Institute, King’s College London, London, UK; USC Norris Comprehensive Cancer Center, Epigenetics and Gene Regulation, Los Angeles, CA 90089, USA; Molecular and Computational Biology Department, USC Dornsife College of Letters, Arts and Sciences, Los Angeles, CA 90089; USC Stem Cell Initiative, Los Angeles, CA 90089, USA

## Abstract

Neutrophils are the most abundant white blood cells in humans and constitute one of the first lines of defense in the innate immune response. Neutrophils are extremely short-lived cells, which survive less than a day after reaching terminal differentiation. Thus, little is known about how organismal aging, rather than the daily cellular aging process, may impact neutrophil biology. In addition, accumulating evidence suggests that both immunity and organismal aging are sex-dimorphic. Here, we describe a multi-omic resource of mouse primary bone marrow neutrophils from young and old female and male mice, at the transcriptomic, metabolomic and lipidomic levels. Importantly, we identify widespread age-related and sex-dimorphic regulation of ‘omics’ in neutrophils, specifically regulation of chromatin. Using machine-learning, we identify candidate molecular drivers of age-related and sex-dimorphic transcriptional regulation of neutrophils. We leverage our resource to predict increased levels/release of neutrophil elastase in male mice. To date, this dataset represents the largest multi-omics resource for the study of neutrophils across biological sex and ages. This resource identifies molecular states linked to neutrophil characteristics linked to organismal age or sex, which could be targeted to improve immune responses across individuals.

## Introduction

Neutrophils are the most abundant cell type in human blood, representing 50-70% of leukocytes in humans throughout life (Nah et al., 2018). These cells are continually produced in the bone marrow and released into circulation to participate in immune surveillance (Furze and Rankin, 2008; Shah et al., 2017). Neutrophils are also very short-lived cells, with estimated cellular lifespan of ∼6 hours once released in the bloodstream, in a process that has been dubbed “Neutrophil aging”, distinct from organismal aging (Shah et al., 2017; Zhang et al., 2015). Neutrophils perform a number of key effector functions, including the production of antimicrobial granules and of “neutrophil extracellular traps” [NETs] (Shah et al., 2017; Sollberger et al., 2018). Although neutrophils are essential for the defense against infections as the “first line of defense”, their aberrant activation can also aggravate inflammatory disease (Furze and Rankin, 2008; Shah et al., 2017). Indeed, emerging evidence suggests that neutrophils play important roles in chronic inflammation (Soehnlein et al., 2017).

Mammalian aging is characterized by systematic immune dysfunction and chronic sterile inflammation, a phenomenon dubbed “inflamm-aging” (Franceschi and Campisi, 2014). Due to the short-lived nature of neutrophils, they have not yet been extensively studied throughout <u>organismal</u> aging, despite emerging evidence suggesting that neutrophils from aged organisms may be partially dysfunctional (Tseng and Liu, 2014). Indeed, neutrophil dysfunction in aging organisms may help drive aspects of age-related inflammatory phenotypes, including the neutrophils’ ability to resolve inflammation, to undergo NETosis and to secrete proteases from their intracellular granules (Sapey et al., 2014; Tseng and Liu, 2014). Although changes in gene expression regulation throughout lifespan have been reported across many tissues and cell types (Benayoun et al., 2019; Lai et al., 2019), how organismal aging (rather than “daily” cellular aging) affects neutrophils remains largely unknown.

Females and males differ in many aspects of innate and adaptive immune responses, which has been proposed to underlie lifelong health disparities between sexes (Klein and Flanagan, 2016). Interestingly, many such immune differences could stem from differential “omic” regulation across a number of immune cell types (Gal-Oz et al., 2019; Marquez et al., 2020; Qu et al., 2015). Although transcriptional differences between female and male murine neutrophils have started to be profiled by the ImmGen consortium (Gal-Oz et al., 2019), how these differences interplay with organismal aging, and whether they are accompanied by other phenotypical differences in neutrophil biology is still largely unclear. Accumulated studies suggest that aspects of neutrophil biology are sex-dimorphic, for instance in their ability to produce lipid-derived inflammatory mediators (Soehnlein et al., 2017), or their functional modulation by testosterone (Markman et al., 2020). However, the potential biological and genomic pathway underlying sex-dimorphism in neutrophils, as well as the extent of sex-dimorphism in neutrophil biology, are still largely unexplored.

To gain insights into how neutrophils are regulated throughout organismal lifespan and as a function of biological sex, we generated a “multi-omic” resource covering transcriptome, metabolome and lipidome profiling of primary bone marrow mouse neutrophils. We identified widespread age-related and sex-dimorphic omic regulation, including transcriptional regulation of chromatin-related gene expression. Using ATAC-seq, we showed that transcriptional remodeling of chromatin-related pathways was associated with overall differences in the chromatin architecture of neutrophils isolated from young *vs.* old and female *vs.* male mice. Machine-learning showed that specific transcription factors could help predict age-related and sex-dimorphic gene regulation in neutrophils, thus identifying potential molecular drivers of phenotypic diversity with organismal age and biological sex. Finally, we leveraged our “omic” resource and were able to predict sex-differences in serum levels of neutrophil primary granule-derived elastase in control and sepsis-like conditions.

## Results

### A transcriptomic, metabolomic and lipidomic atlas of primary mouse bone marrow neutrophils with respect to aging and sex

To understand how neutrophils may be regulated during aging and as a function of sex, we obtained primary bone marrow neutrophils from young adult (4-5 months) and old (20-21 months) C57BL/6Nia female and male mice, with 4 animals in each group (**Figure 1A**). Primary neutrophils were isolated from the bone marrow of long-bones using magnetic-activated cell sorting (MACS). Multi-omic profiling was then performed on purified neutrophils: (i) transcriptome profiling by RNA-sequencing [RNA-seq], (ii) metabolite profiling by untargeted liquid chromatography coupled with mass spectrometry (LC-MS) and (iii) lipid profiling by targeted MS (**Figure 1A**).

**Figure 1:**
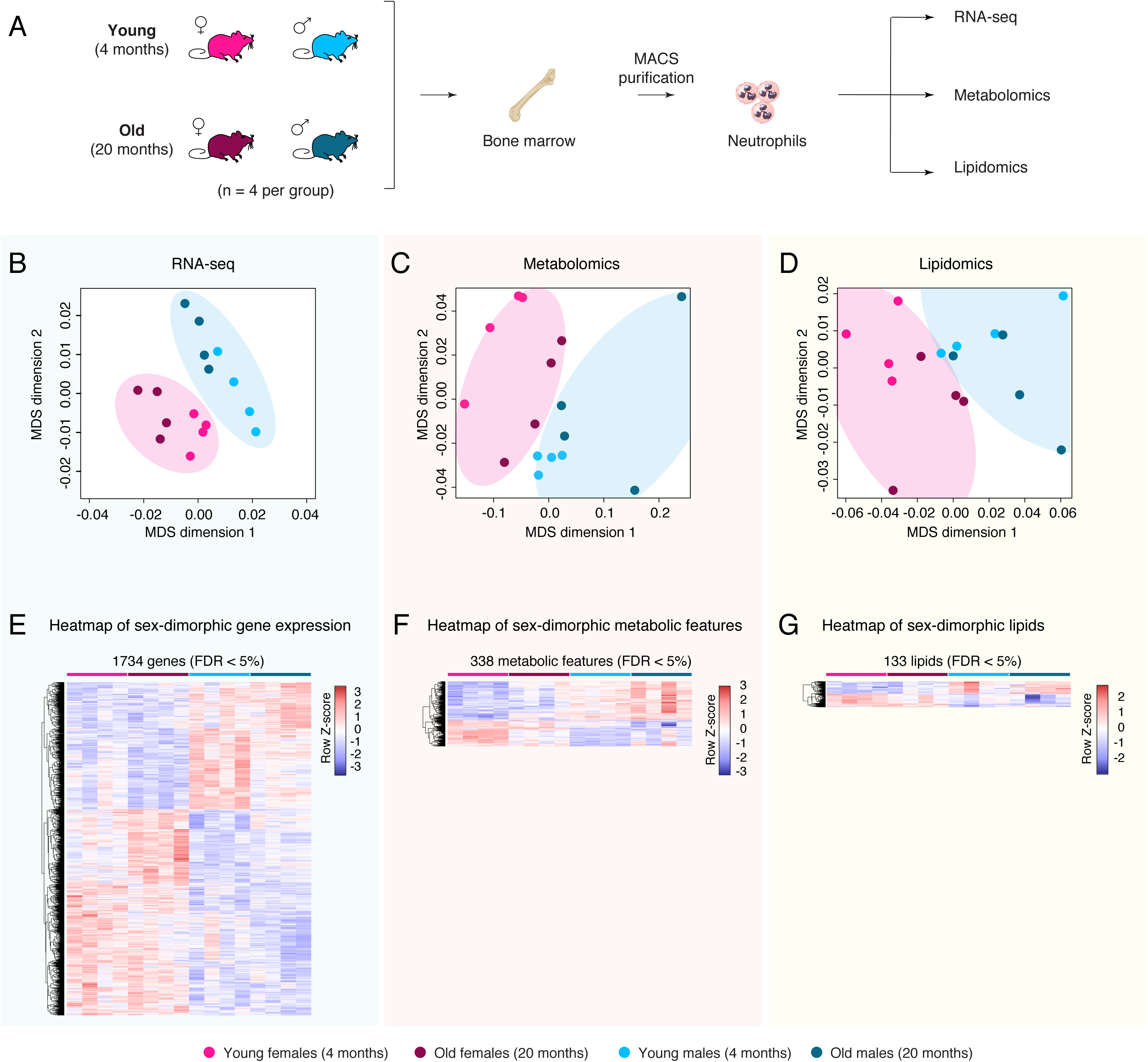
A multi-omic analysis of primary mouse bone marrow neutrophils during aging and with respect to sex. (**A**) Experimental setup scheme. (**B-D**) Multidimensional Scaling analysis results of RNA expression by RNA-seq (B), untargeted metabolomics (C), or targeted lipidomics (D). (**E-G**) Heatmap of significant (FDR < 5%) sex-dimorphic genes (E), metabolic features (F) or lipid species (G). Significance of gene regulation by RNA-seq was calculated by DESeq2, and significance of metabolic features or lipid species regulation were calculated by limma.

As a first-level analysis to evaluate the similarity of our ‘omics’ datasets, we utilized multidimensional scaling [MDS] (Chen and Meltzer, 2005). MDS analysis for RNA-seq, metabolomics and lipidomics datasets showed clear separation of samples by the sex of the animals, regardless of age (**Figure 1B-D**). In contrast, although young and old samples seemed to separate within each sex for each omics, <u>global</u> separation by age regardless of sex was not as clearly observed as for sex (**Figure 1B-D**). To better understand the nature of molecular differences between neutrophils from young *vs.* old [with sex as a covariate], and female *vs.* male animals [with age as a covariate], we identified transcriptional, metabolic and lipidomic features with significant age- or sex-related differential regulation at a False Discovery Rate [FDR] < 5% using multivariate linear modeling (see methods; **Figure 1E-G** and **S1A-F**; **Supplementary Table S1A-F**).

Consistent with the MDS analysis, we were able to identify genes, metabolites and lipid species with significant differential regulation with respect to biological sex (FDR < 5%; **Figure 1E-G** and **S1A-D**; **Supplementary Table S1B,D,E**). Genes that were differentially expressed with respect to sex were located throughout the genome, without a clear enrichment of genes with sex-biased expression on sex chromosomes, suggesting that sex-dimorphic gene expression in neutrophils is not the mere byproduct on genomic organization (**Figure S1A,E**). Importantly, there were no apparent biases in the purity of MACS-purified neutrophils from female *vs.* male animals (**Figure S2A-C**), and no significant difference in starting neutrophil abundance from bone-marrow of female *vs.* male animals (**Figure S2D**). Thus, it is unlikely that the prevalence of sex-dimorphic molecular phenotypes of bone-marrow neutrophils is the mere result of a systematic purification bias between sexes.

Although aging was associated with clear changes in neutrophil gene expression profiles as detected by RNA-seq (**Figure S1A,F**), only one metabolic feature was significantly changed with aging, and no significant lipidomic changes were observed (**Figure S1A,F**; **Supplementary Table S1C,F**). This is consistent with the overall poorer age-driven separation observed in the MDS analysis for metabolomics and lipidomics (**Figure 1B-C**). Higher biological variability in metabolites and lipids might explain the lack of widespread significant changes with age. Alternatively, it is possible that the paucity of age-related metabolic and lipidomic differences may stem from biological specificities of neutrophil (*e.g.* their short cellular lifespan). When female and male neutrophils were analyzed separately, we found that the transcriptional impact of aging on neutrophils was very strongly correlated between sexes (Rho = 0.593 and p ∼ 0 in significance of correlation test; **Figure S1G**). Thus, although female and male neutrophils show clear differences throughout life, the trajectories of neutrophils with organismal aging are highly similar regardless of sex. To note, we observed a greater amount of significant transcriptional changes with organismal age in male neutrophils compared to female neutrophils (**Figure S1H**), suggesting that male neutrophils may be more susceptible to aging expression changes than female neutrophils. Together, our results indicate that bone marrow neutrophils are highly sex-dimorphic at the molecular level, and that these sex differences persist (or may be amplified) with organismal aging.

### Functional enrichment analysis of neutrophil ‘omics’ reveals age-related regulation of chromatin-related pathways

We first focused on the impact of organismal aging on neutrophils function using pathway enrichment analysis (**Figure 2A-C** and **S3A-B; Supplementary Table S2**). To note, we only analyzed age-related functional remodeling <u>at the transcriptomic level</u>, since metabolomic and lipidomic profiles showed few to no significant changes with aging (**Supplementary Figure S1A**).

First, we assessed how coherent and coordinated age-related changes are by performing a network analysis using NetworkAnalyst 3.0 (Zhou et al., 2019) (**Figure 2A**). The network was constructed using significantly age-regulated genes and protein-protein interaction [PPI] information derived from IMEx/InnateDB data (Breuer et al., 2013), a knowledgebase specifically geared for analyses of innate immune networks. Our analysis revealed that significantly age-regulated genes are part of a clearly interconnected network (**Figure 2A**), suggesting coordinated (*i.e.* non-random) shifts in the neutrophil functional landscape with organismal aging.

**Figure 2:**
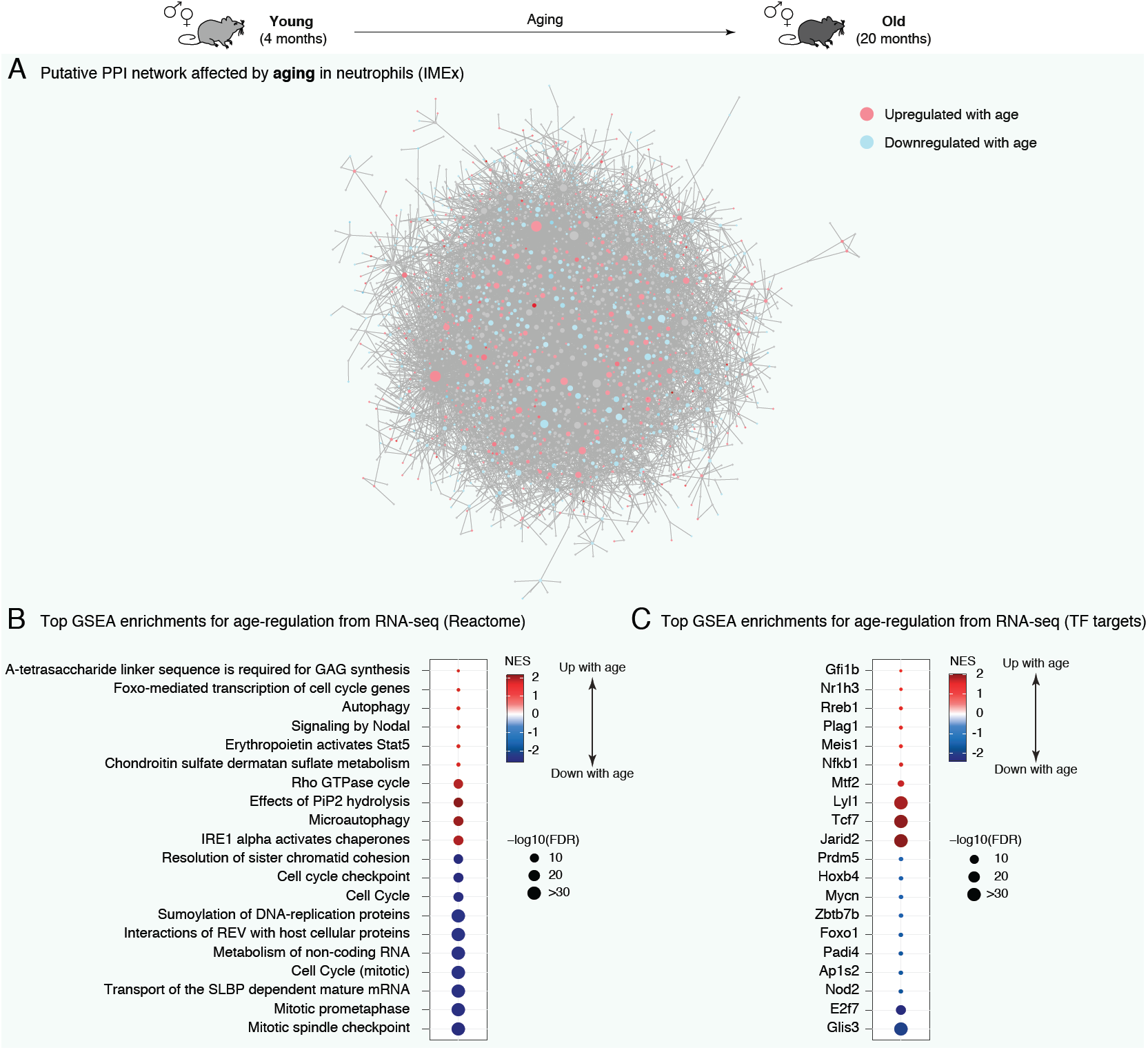
Misregulated pathways in bone marrow neutrophils during aging reveals downregulation of chromatin-related pathways. (**A**) NetworkAnalyst predicted PPI network for significant age-regulated genes in neutrophils. Network edges are based on IMEx/InnateDB data, a knowledgebase specifically geared for analyses of innate immune networks (Breuer et al., 2013). Blue (red) shades are associated to decreased (increased) gene expression during aging. (**B-C**) Top enriched gene sets from REACTOME (B) and transcription factor targets (C) using GSEA for differential RNA expression. Only the top 10 most up- and top 10 most downregulated gene sets are plotted for readability. Full lists and statistics available in **Supplementary Table S2**. Also see **Supplementary Figure S3**. All shown pathways and genes such as FDR < 5%. NES: Normalized Enrichment Score (for GSEA analysis). FDR: False Discovery Rate.

We then asked which specific pathways were misregulated with organismal aging in bone marrow neutrophils (**Figure 2B** and **S3A,B; Supplementary Table S2**). Some immunity or antigen processing-related linked pathways were significantly upregulated with organismal aging across functional annotations sources by Gene Set Enrichment Analysis [GSEA] (*e.g*. REACTOME “EPO activates Stat5”, Gene Ontology “peptide-antigen binding”, KEGG “primary immunodeficiency”, etc.; **Figure 2B** and **S3A,B**; **Supplementary Table S2**). The upregulation of immune-related pathways was also observed with Ingenuity Pathway Analysis [IPA] (**Supplemental Table S2E**). This suggests that neutrophil-associated immunity in older animals could have different characteristics, consistent with the widespread observations of age-related immune dysfunction or “inflamm-aging” (Franceschi and Campisi, 2014).

Intriguingly, the top downregulated pathways with organismal aging were overwhelmingly related to chromatin and cell cycle regulation (*e.g*. REACTOME “Cell cycle”, Gene Ontology “chromosome”, KEGG “DNA replication”, etc.) (**Figure 2B** and **S3A,B; Supplementary Table S2**). The downregulation of chromatin and cell cycle-related pathways was also observed with IPA (**Supplemental Table S2E**). Specifically, the expression of 18 histone-encoding genes were significantly downregulated with aging (**Supplemental Table S1A**). Changes in the global regulation of chromatin components are especially relevant to neutrophil biology. Indeed, the overall chromatin organization of neutrophils differs from that of other cells (Chen et al., 2016), including increased chromatin compaction or “supercontraction” (Denholtz et al., 2020; Zhu et al., 2017). In addition, neutrophils can extrude their chromatin to produce Neutrophil Extracellular Traps [NETs] in response to pathogens (Papayannopoulos, 2018). The extrusion of chromatin directly participates in pathogen killing, notably thanks to the antimicrobial properties of histone proteins (Brinkmann et al., 2004; Brinkmann and Zychlinsky, 2012). In addition, neutrophil chromatin relaxation can also be potentiated by nuclear-translocated granule-derived proteases (*e.g.* Elane, Mpo), which ultimately help in bacterial killing (Papayannopoulos et al., 2010). Finally, although neutrophils are fully post-mitotic, cell-cycle genes play important roles in controlling the production of NETs (Amulic et al., 2017). Thus, our analysis suggests that neutrophils may experience global changes in chromatin organization with organismal aging, a feature that may impact their ability to respond to infectious challenges.

We also observed that several autophagy-related pathways were misregulated in old neutrophils (*e.g*. REACTOME “Autophagy” and “Microautophagy”, Gene Ontology “positive regulation of autophagosome assembly”, IPA “Autophagy” and “mTOR Signaling”, etc.) (**Figure 2B** and **Supplementary Table S2B,C,E**). This is consistent with misregulation of autophagy pathways being a “hallmark of aging” (Lopez-Otin et al., 2013; Wong et al., 2020). More specifically for neutrophils, control of autophagy is critical for normal neutrophil differentiation (Riffelmacher et al., 2017), regulation of NET formation (Park et al., 2017) and regulation of neutrophil degranulation (Bhattacharya et al., 2015). Interestingly, neutrophils from older humans show evidence of increased degranulation compared to those from young donors (Sapey et al., 2014).

To identify candidate upstream regulators with age-related changes in regulatory activity, we investigated whether target genes of transcription factors [TFs] were misregulated with aging (**Figure 2C; Supplementary Table S2D**). We obtained lists of TF target genes derived from ChEA, JASPAR or GEO, based on gene sets from the Harmonizome knowledgebase (see methods) (Rouillard et al., 2016). Importantly, we restricted our testing to TFs with detectable expression in neutrophils according to our RNA-seq dataset. To note, the “TF” target set from GEO obtained from Harmonizome (Rouillard et al., 2016) includes several non-TF regulators, such as chromatin-remodeling enzymes or signaling receptors, which for simplicity are still referred to hereafter as TFs. Consistent with functional pathway enrichment results, we observed significant age-related downregulation of the target genes of E2f7, which is known to regulate cycle-related gene expression (Grant et al., 2013; Mitxelena et al., 2018; Yuan et al., 2019) (**Figure 2C**). Although E2f7 has not been studied in depth in neutrophils, it is highly expressed in committed neutrophil progenitors (Kim et al., 2017). Targets of Foxo1 were also significantly downregulated with aging in neutrophils (**Figure 2C**). Foxo TFs are the primary targets of the aging-regulating insulin/insulin-like growth factor 1 signaling (Brown and Webb, 2018), and Foxo1 regulates neutrophil-mediated bacterial immunity (Dong et al., 2017). Finally, known targets of Padi4 were predicted to be significantly downregulated with aging (**Figure 2C**). Padi4 is a peptidylarginine deaminase, which catalyzes histone citrullination, and plays a key role in NETosis *in vivo* (Li et al., 2010a; Rohrbach et al., 2012; Thiam et al., 2020). Transcript levels of *Padi4* were significantly upregulated with aging (FDR = 6.9×10^-3^; **Figure S3C**; **Supplementary Table S1B**). This is consistent with reports of histone citrullination associating to transcriptional repression (Cuthbert et al., 2004; Denis et al., 2009; Li et al., 2010b; Li et al., 2008; Wang et al., 2012; Wang et al., 2004), although the directionality of transcriptional regulation may depend on cofactors and cell types (Christophorou et al., 2014). Together, our functional pathway analysis suggests that major aspects of chromatin metabolism are shifting in aging primary bone marrow neutrophils.

### Functional enrichment analysis of neutrophil ‘omics’ reveals widespread sex-dimorphism in chromatin-related pathways

We next analyzed the functional impact of sex-dimorphic regulation in neutrophils at the transcriptomic, metabolomic and lipidomic levels (**Figure 3A-F** and **S4A-C**; **Supplementary Table S3**). As for aging, we first assessed the interconnection of sex-dimorphic genes using NetworkAnalyst 3.0 to reconstruct putative PPI networks, based on IMEx/InnateDB interactions (Breuer et al., 2013), for female *vs.* male-biased gene expression (**Figure 3A,B**). Sex-biased gene programs showed clear interconnection, again suggesting coherent differences with likely functional impact (**Figure 3A,B**). The top connected node in the female-biased gene network, *Iqcb1* (also known as *Nphp5*), encodes a primary cilia component and has been linked to nephronophthisis, a severe kidney disease (**Figure 3A**) (Hossain et al., 2020; Srivastava et al., 2017). Interestingly, *Irf8* was the top most connected node in the male-biased gene network (**Figure 3B**). Although it has not been specifically studied in neutrophils, Irf8 is a TF-related to interferon signaling that plays a key role in myeloid cell fate determination (usually suppressing neutrophil differentiation) and antigen-presentation regulation (Marquis et al., 2011; Papayannopoulos, 2018; Yáñez et al., 2015).

**Figure 3:**
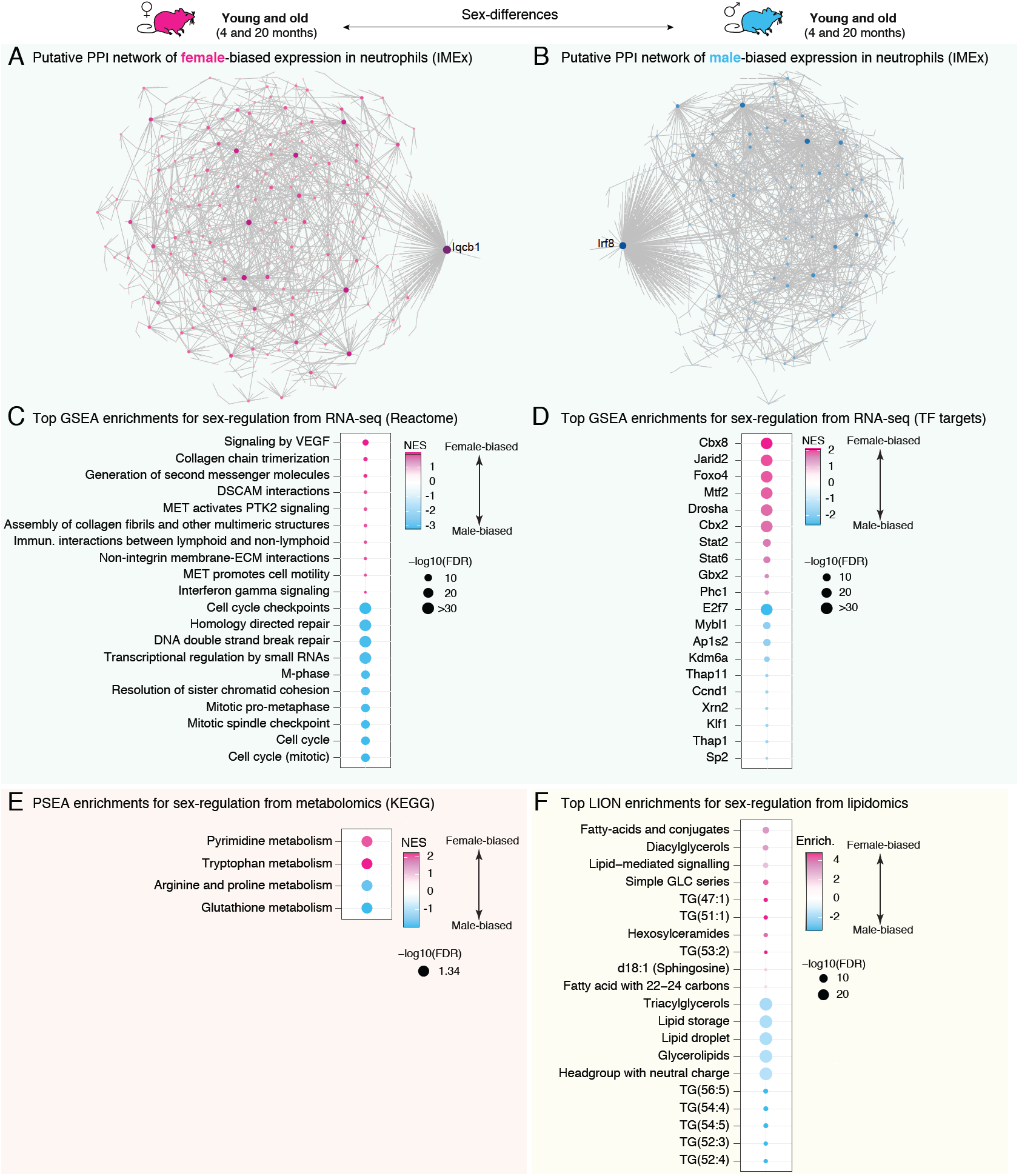
Sex-dimorphic pathways in bone marrow neutrophils reveals differential regulation of chromatin-related pathways. (**A-B**) NetworkAnalyst predicted PPI network for genes displaying significant bias in gene expression by RNA-seq towards female (A) or male (B) neutrophils. Network edges are based on IMEx/InnateDB data, a knowledgebase specifically geared for analyses of innate immune networks (Breuer et al., 2013). (**C-D**) Top enriched gene sets from REACTOME (C) and transcription factor targets (D) using GSEA for differential RNA expression. (**E**) Phenotype Set Enrichment Analysis (PSEA) for differential metabolite regulation. (**F**) Top Lipid Ontology (LION) functional enrichment analysis for differential lipid regulation. For (C-F), the top 10 most up- and top 10 most downregulated gene sets (if that many) are plotted for readability. Full lists and statistics available in **Supplementary Table S3**. Also see **Supplementary Figure S4**. All shown pathways and genes such as FDR < 5%. NES: Normalized Enrichment Score (for GSEA and PSEA analysis). FDR: False Discovery Rate.

Next, we asked which functional pathways were regulated in a sex-dimorphic fashion in neutrophils (**Figure 3C** and **S4A,B**; **Supplementary Table S3**). Functional enrichment analysis revealed that significant female-biased pathways encompassed a number of extra-cellular matrix [ECM] and cell surface related pathways (*e.g*. REACTOME “collagen chain trimerization”, Gene Ontology “external side of plasma membrane”, KEGG “cell adhesion molecules”, etc.), according to GSEA and IPA (**Figure 3C** and **S4A,B**; **Supplementary Table S3A-D**). Indeed, collagen-encoding genes *Col1a1* and *Col1a2* were expressed at significantly higher levels in female neutrophils (**Supplementary Table S1B**). Collagen production, usually from fibroblasts, is a key event in tissue fibrosis (Wynn, 2008), and neutrophils play an important role in the development of fibrotic lesions (Frangou et al., 2019; Gregory et al., 2015; Koyama and Brenner, 2017). Alternatively, differential expression of cell surface-related genes may also lead to differences in the ability of neutrophils to migrate through endothelial barriers (Muller, 2016). Interestingly, a number of autophagy-related gene sets were female-biased in our dataset according to GSEA (*e.g*. Gene Ontology “macroautophagy”, “autophagy”, or “autophagosome membrane”, KEGG “Regulation of autophagy”) (**Supplementary Table S3A,C**). IPA also identified “autophagy”-related genes to be significantly enriched among sex-dimorphic genes (**Supplementary Table S3E**). As mentioned above, autophagy control is critical for neutrophil biology (Park et al., 2017; Riffelmacher et al., 2017), including the regulation of neutrophil degranulation (Bhattacharya et al., 2015). Intriguingly, chromatin- and cell cycle-related pathways were found to be overwhelmingly biased for higher expression in male neutrophils (*e.g*. REACTOME “Cell cycle”, Gene Ontology “chromosome”, KEGG “DNA replication”, etc.), according to GSEA and IPA (**Figure 3C** and **S4A,B**; **Supplementary Table S3**). Indeed, 21 histone-encoding genes showed significantly higher expression in male *vs.* female neutrophils (**Supplementary Table S1B**). The sex-dimorphic regulation of chromatin and cell cycle-related pathways suggests that male and female neutrophils have differences in their overall chromatin metabolism, and thus overall chromatin organization.

We also investigated whether sex-dimorphic gene expression was correlated with differential predicted activity of TFs expressed in neutrophils (**Figure 3D; Supplementary Table S3D**). Consistent with pathway analysis results, targets of cell cycle-related TFs E2f7 and Mybl1 (also known as a-Myb) (Oh and Reddy, 1999; Yuan et al., 2019) were more highly expressed in male neutrophils (**Figure 3D**). Female neutrophils showed higher expression of targets of Foxo4 (**Figure 3D**). Interestingly, the insulin Insulin/Igf1 signaling pathway, which regulates Foxo TF activity, has been linked to multiple sex-dimorphic phenotypes (*e.g.* longevity) (Austad and Bartke, 2015; Penniman et al., 2019).

We next examined sex-dimorphism in metabolic pathways based on our metabolomics data and identified a number of metabolic pathways with significant sex-dimorphic patterns (**Figures 3E** and **S4C; Supplementary Table S3F**). Interestingly, metabolic pathways related to nucleotide metabolism were differentially regulated between females *vs.* male neutrophils (*e.g.* KEGG “Pyrimidine Metabolism”; **Figures 3E** and **S4C; Supplementary Table S3F**). Indeed, a number of sex-dimorphic metabolic peaks were predicted to represent nucleotide species (*e.g.* AMP, ATP, GMP; manually validated nucleotide species in **Supplementary Table S1D**). Sex-differences in nucleotide pools are consistent with our transcriptomic observations of sex-dimorphism in the predicted activity of (i) pathways directly related to nucleotide metabolism (*e.g.* KEGG “pyrimidine metabolism”, Reactome “metabolism of nucleotides”, GO “purine nucleobase biosynthetic process”), which could lead to direct changes in the composition of nucleotide pools, and (ii) pathways responsive to nucleotide levels, such as signaling by Rho-GTPases (*e.g.* Reactome “signaling by Rho GTPases”, GO “GTPase activator activity”), which can regulate neutrophil recruitment (Baker et al., 2016) (**Supplementary Table S3A-C,E**). Among nucleotides, the impact of adenine derivatives in neutrophils has been studied more extensively. ATP/ADP levels are regulated in neutrophils as a function of activation state and ‘cellular age’ (Richer et al., 2018). Adenine nucleotides derived from neutrophils have been suggested to have anti-inflammatory effects (Cadieux et al., 2005; Eltzschig et al., 2004; Hasko and Cronstein, 2013), increase endothelial barrier function, attenuate neutrophil adhesion to endothelial cells, and thus modulate transendothelial migration (Eltzschig et al., 2006). Functional analysis of our metabolomic data also suggested significant sex-differences in amino-acid metabolism pathways, such as tryptophan and arginine/proline metabolism (**Figures 3E** and **S4C**). Interestingly, arginine and tryptophan metabolism are known to play important immunoregulatory roles in neutrophils (Mondanelli et al., 2019). However, the final effect of differential metabolite levels belonging to these pathways between sexes will need further investigation.

We then used the Lipid Ontology [LION] framework (Molenaar et al., 2019) to analyze sex-dimorphism in lipid functions based on our lipidomics data (**Figures 3F, S1D** and **Supplemental Table S3H**). Interestingly, male neutrophils were strongly enriched for triacylglycerols [TAG], which are stored in lipid droplets, according to the LION analysis (**Figures 3F** and **Supplemental Table S3G**). This was also independently observed when analyzing lipidomics data by class (**Figure S1D**). In contrast, female neutrophils were enriched for diacylglycerols [DAG] (**Figures 3F** and **S1D**). Consistently, genes associated to the Gene Ontology term “negative regulation of lipid storage” (GO:0010888) were expressed at significantly higher levels in female neutrophils according to our transcriptomic data (FDR = 4.33×10^-3^; **Supplemental Table S3C**). Lipid droplets are very important in the control of neutrophil differentiation and as a source for inflammatory mediators (Jarc and Petan, 2020; Riffelmacher et al., 2017). Adipose triglyceride lipase (Atgl), encoded by the *Pnpla2* gene, is known to be a key regulator of neutral lipid storage into lipid droplets in neutrophils (Schlager et al., 2015). Interestingly, *Pnpla2* is expressed at higher RNA levels in female bone marrow neutrophils (FDR = 0.01; **Figure S4D**; **Supplemental Table S1B**). Indeed, Atgl catalyzes the conversion of TAG to DAG, consistent with higher levels of DAG in female neutrophils and of TAG in male neutrophils (**Figure 3F** and **S1D**). Lower levels of *Pnpla2*, thus mimicking a more “male” state, can lead to abnormal neutral lipid/triglycerides accumulation in neutrophils, increased chemotactic ability, and reduced release of proinflammatory lipid mediators (Schlager et al., 2015). In addition, *Atg7* is crucial for generation of free fatty acids from lipid-droplets in neutrophils, and *Atg7*-deficient neutrophils show increased lipid-droplet storage (Riffelmacher et al., 2017). Consistently, *Atg7* is expressed at higher RNA levels in female neutrophils (FDR = 0.03; **Figure S4D**; **Supplemental Table S1B**). Finally, *Lpin1* is crucial for the synthesis of DAG from phosphatidic acid, and is known to play a role in the regulation of macrophage inflammation (Meana et al., 2014). Consistent with observed increased DAG levels, *Lpin1* is also expressed at higher RNA levels in female neutrophils (FDR = 0.03; **Figure S4D**; **Supplemental Table S1B**). Thus, both lipidomics and transcriptomics data suggest that male neutrophils have increased neutral lipid storage (or decreased usage) compared to female neutrophils (**Figure S4E**). Because of the functional importance of lipid droplets in neutrophils biology, these observations are likely to have deep functional consequences on overall neutrophil function between sexes.

Finally, we performed an integrated pathway analysis using the IMPaLA framework (Kamburov et al., 2011), which can combine information from transcriptomics and metabolomics/lipidomics analyses (**Supplementary Table S3H**). We observed an extremely strong joint enrichment for male-biased molecules to be in pathways linked to cell cycle, DNA metabolism and chromatin-related pathways (*e.g.* “Cell Cycle”, “DNA Replication”, “Chromosome Maintenance”; **Supplementary Table S3H**), further supporting the notion that neutrophil chromatin architecture may be regulated in a sex-dimorphic manner. In contrast, female-biased enriched pathways were enriched for lipid-metabolism related pathways (*e.g.* “phospholipases”, “triacylglycerol degradation”, “phospholipid metabolism”; **Supplementary Table S3H**). These joint observations provide an independent and integrated confirmation of our observations for each individual “omic” layer. Together, our multi-omic analyses suggest that several major aspects of neutrophil functions are regulated in a sex-dimorphic fashion and may thus ultimately underlie sex-differences in immune responses.

### The global chromatin architecture of neutrophils is modulated by sex and organismal age

Our functional enrichment analyses revealed that chromatin-related pathways were significantly modulated by both sex and organismal age in primary bone marrow neutrophils, which is expected to result in profound changes in overall chromatin architecture. Neutrophil hold their chromatin in a compacted polylobular nuclear architecture, which has earned them the name of “polymorphonuclear” cells (Chen et al., 2016; Denholtz et al., 2020; Zhu et al., 2017). More than just a gene expression regulatory layer, neutrophil chromatin also directly participates in antimicrobial responses as a key component of NETs (Brinkmann et al., 2004; Brinkmann and Zychlinsky, 2012; Papayannopoulos, 2018). Thus, because of the unique functions and regulation of neutrophil chromatin, such regulatory differences are expected to lead to profound changes in neutrophil biology. Thus, to directly evaluate changes in local and overall chromatin accessibility and structure, we utilized the Assay for Transposon-Accessible Chromatin followed by sequencing (ATAC-seq) (Buenrostro et al., 2013; Corces et al., 2017). ATAC-seq takes advantage of adapter-loaded Tn5 particles inserting into accessible regions of the genome to reveal the underlying chromatin landscape, yielding information in the same experiment about both local chromatin accessibility (Buenrostro et al., 2013) and higher order chromatin organization (Fortin and Hansen, 2015; Schep et al., 2015).

To uncover potential differences in the neutrophil chromatin landscape, we isolated primary bone marrow neutrophils from a new independent cohort of young adult (4-5 months) and old (20-21 months) C57BL/6Nia female and male mice and performed ATAC-seq profiling on freshly isolated cells (**Figure 4A** and **S5A-G**). In contrast with our transcriptomic, metabolomic and lipidomic data, MDS analysis on ATAC-seq peaks showed that aging was a better factor of sample separation than sex <u>when examining only local differences</u> in chromatin accessibility (**Figure S5C**). In addition, there were substantially more regions with age-related compared to sex-dimorphic differential accessibility (**Figure S5D-G**), consistent with our observations in terms of numbers of differentially expressed genes with respect to age and sex (**Figure S1A**).

**Figure 4:**
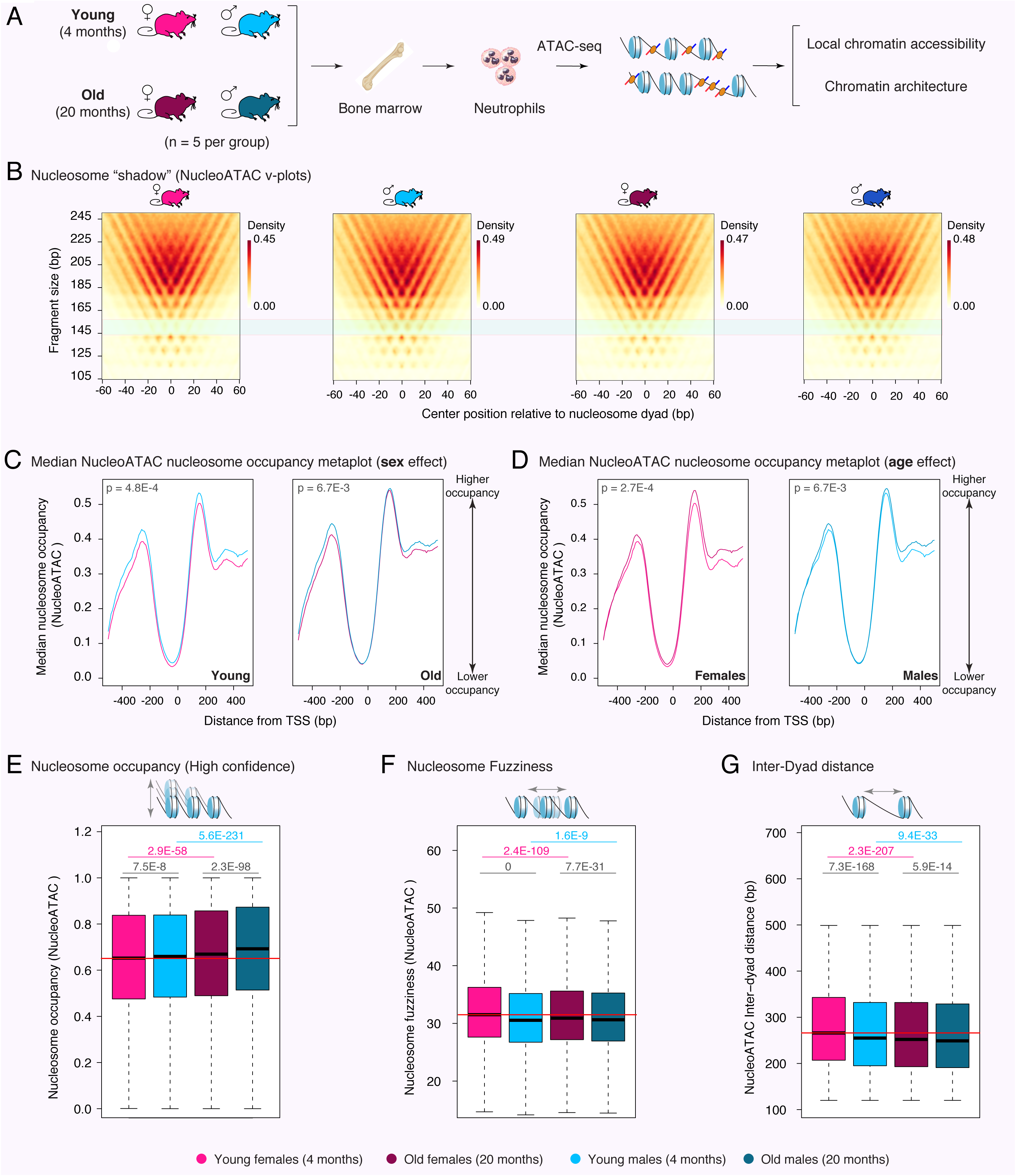
ATAC-seq analysis reveals age- and sex-related differences in the chromatin architecture of bone marrow neutrophils. (**A**) Setup scheme for ATAC-seq experiment. (**B**) NucleoATAC v-plots. The light green box overlay reveals a v-plot region where males have higher signal than females, regardless of age, suggesting differences in nucleosomal architecture. (**C-D**) Metaplot analysis of median nucleosome occupancy (as calculated by NucleoATAC) around the TSS of neutrophil-expressed genes, to analyze sex- (C) or age-related (D) differences in neutrophil nucleosome occupancy. Note that increased occupancy is observed in male compared to female neutrophils, as well as in old compared to young neutrophils. Significance of the difference between median occupancy profiles at TSSs assessed using a Kolmogorov-Smirnov goodnest-of-fit test. (**E**) Boxplot of nucleosome occupancy as calculated by NucleoATAC for high-confidence nucleosomes across groups. Also see **Supplementary Figure S5H**. (**F**) Boxplot of nucleosome fuzziness as calculated by NucleoATAC for high-confidence nucleosomes across experimental groups. (**G**) Boxplot of nucleosome inter-dyad distance as calculated by NucleoATAC across experimental groups. The horizontal red line in panels (**E-G**) shows the median value in neutrophils from young females for ease of comparison. Significance in non-parametric Wilcoxon rank-sum tests are reported for panels (**E-G**). Black p-values represent differences between male *vs.* female neutrophils in each age group; pink (blue) p-values represent age-related differences in female (male) neutrophils. Also see **Supplementary Figure S5H,I**.

Since our functional enrichment analyses suggested that overall aspects of chromatin may be differentially regulated with organismal age and sex, we leveraged the NucleoATAC algorithm to analyze the chromatin architecture of neutrophils across conditions (Schep et al., 2015). Interestingly, nucleosome read “v-plots” generated by NucleoATAC revealed a region around fragment size 145-150bp which captured more reads in male *vs.* female neutrophils, regardless of animal age, suggesting global differences in overall nucleosome packaging (**Figure 4B**). To further elucidate differences in overall chromatin architecture, we leveraged three indicators linked to chromatin compaction and organization around nucleosomes: (i) <u>nucleosome occupancy</u>, which indicates how frequently a position is occupied by a nucleosome across a cell population, where higher occupancy is associated to tighter chromatin structure, (ii) <u>nucleosome fuzziness</u>, which indicates how well positioned the nucleosome is, where fuzzier nucleosomes are linked to looser chromatin, (iii) <u>inter-dyad distance</u>, which measure the length of DNA between neighboring nucleosomes, thus providing information about local chromatin compaction, with increased distance associated to less compact chromatin. Nucleosome occupancy metaplots around the transcriptional start sites [TSS] of expressed genes revealed that, in both the young and old groups, male neutrophils showed higher median nucleosomal occupancy compared to female neutrophils (**Figure 4C**). Similarly, regardless of sex, aging was associated to increased median nucleosomal occupancy at the TSS of expressed genes (**Figure 4D**). More generally at the genome-wide level, we observed that male neutrophils had overall higher nucleosomal occupancy compared to female neutrophils, and that median nucleosomal occupancy increased with age (both for high confidence and all nucleosome calls; **Figure 4E, S5H**). We also observed decreased nucleosome fuzziness in male compared to female neutrophils, as well as old compared to young neutrophils (**Figure 4F**). Finally, we observed shorter inter-dyad distance in male *vs.* female, as well as old *vs.* young neutrophils (**Figure 4G**). Together, small but consistent increases in occupancy, decreases in fuzziness and decreases in inter-dyad distance are indicative of an overall more compact chromatin architecture in male *vs.* female and old *vs.* young neutrophils. Finally, when analyzing ATAC-seq fragments at accessible regions (including subnucleosomal fragments), we observed lower median fragment length in male and old neutrophils (**Figure S5I**). While this observation may seem contradictory, overall shorter ATAC-seq fragments in the libraries may result from relatively “more accessible” nucleosome-free regions in the context of an overall tighter chromatin structure.

To independently evaluate overall chromatin tightness as a function of age and sex, we decided to leverage our RNA-seq datasets to examine transcription associated to repetitive elements (**Figure S5J-K; Supplementary Table S1G,H**). Indeed, eukaryotic genomes contain large proportions of repetitive elements, including transposons, whose transcription is usually tightly repressed through compaction in constitutive heterochromatin (Janssen et al., 2018). Consistent with a more compact chromatin architecture, male neutrophils showed significantly lower transcription of repetitive elements compared to females (350 elements with female-biased transcription *vs*. 12 with male-biased transcription; **Figure S5J; Supplementary Table S1H**). Similarly, old neutrophils also showed significantly lower transcription of repetitive elements compared to young neutrophils (115 elements downregulated *vs*. 27 upregulated with age; **Figure S5K; Supplementary Table S1G**). Interestingly, chromatin decompaction is a key limiting step of NETosis initiation (Neubert et al., 2018), suggesting that baseline differences in chromatin compaction in neutrophils of different ages and sex may impact the dynamics and timeline of NETosis progression.

### Machine-learning reveals predictive features of age-related and sex-dimorphic gene regulation in neutrophils

To understand whether age-related and sex-dimorphic gene expression can be predicted from differences in genomic sequence, chromatin landscape and/or regulation by specific TFs, we took advantage of machine-learning (**Figure 5A-C, 6A-C; Supplementary Figure S6A-F**). We used seven machine-learning algorithms to learn models discriminating between (i) up- and down-regulated expression in neutrophils with age, and (ii) male- or female-biased neutrophil gene expression (*i.e.* conditional inference Trees [cTree], Linear Discriminant Analysis [LDA], neural networks [NNET], Logistic Regression [LogReg], random forests [RF], support vector machines [SVM], and gradient boosting [GBM]). The models were trained with three types of features (i) information relating to genomic features (*e.g.* GC-richness), (ii) information relating to chromatin accessibility derived from our ATAC-seq experiments (*e.g.* changes in promoter accessibility) and (iii) information relating to TF target genes (*i.e.* same data as used above in our GSEA).

**Figure 5:**
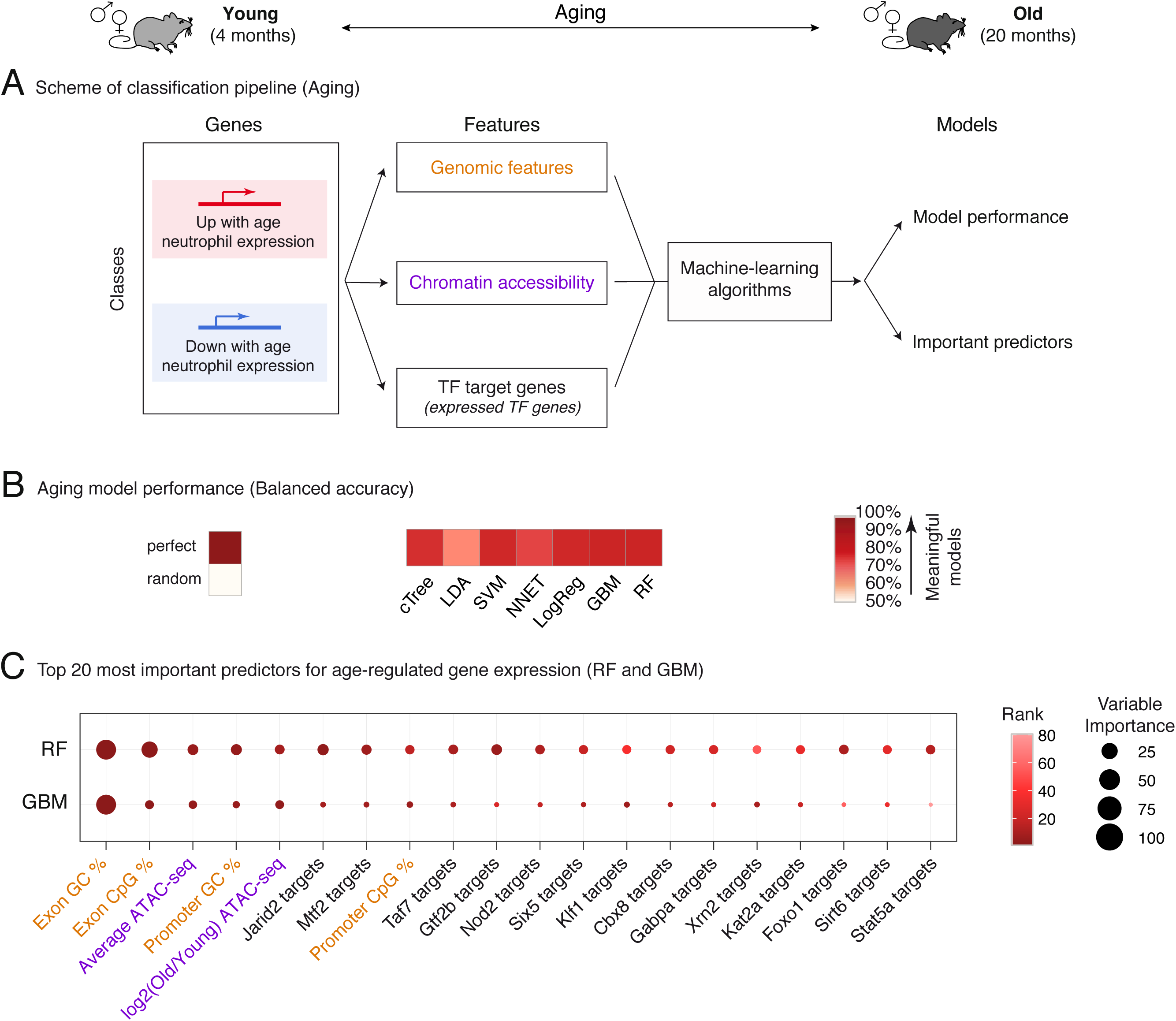
Machine-learning analysis reveals that age-regulated gene expression can be predicted by genomic and epigenomic features. (**A**) Scheme of our machine-learning pipeline for up *vs.* downregulated genes in neutrophils with aging. (**B**) Balanced classification accuracy over different machine-learning model algorithms. The accuracy of the models is measured using held-out testing data. ‘Random’ accuracy illustrates the accuracy of a meaningless model (∼50%), and ‘perfect’ that of a model with no mistakes (100%). All tests were more accurate than random (see other measures of model performance in **Figure S6A-C** and **Supplementary Table S4A**). (**C**) Top 10 most important features from Random Forest models and Gradient Boosting Machine models. A rank product approach was used to determine overall top predictive features from both models. High values for variable importance indicate most influential predictors. cTree: conditional inference tree, LDA: linear discriminant analysis, SVM: support vector machine, NNET: neural network, LogReg: logistic regression, GBM: gradient boosting machine, RF: random forest.

**Figure 6:**
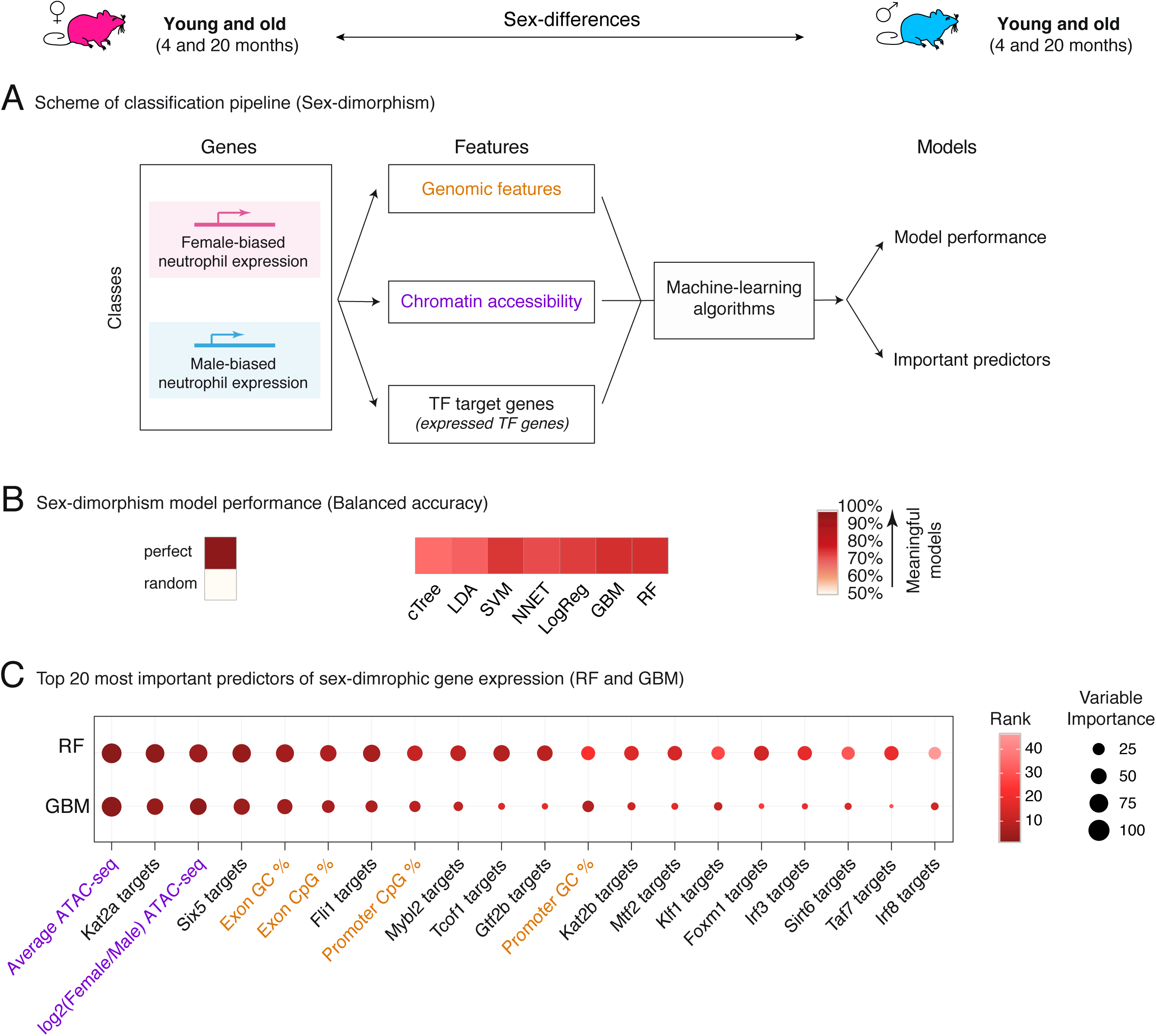
Machine-learning analysis reveals that sex-dimorphic gene expression can be predicted by genomic and epigenomic features. (**A**) Scheme of our machine-learning pipeline for male *vs.* female-biased gene expression in neutrophils. (**B**) Balanced classification accuracy over different machine-learning model algorithms. The accuracy of the models is measured using held-out testing data. ‘Random’ accuracy illustrates the accuracy of a meaningless model (∼50%), and ‘perfect’ that of a model with no mistakes (100%). All tests were more accurate than random (see other measures of model performance in **Figure S6D-F** and **Supplementary Table S4C**). (**C**) Top 10 most important features from Random Forest models and Gradient Boosting Machine models. A rank product approach was used to determine overall top predictive features from both models. High values for variable importance indicate most influential predictors. cTree: conditional inference tree, LDA: linear discriminant analysis, SVM: support vector machine, NNET: neural network, LogReg: logistic regression, GBM: gradient boosting machine, RF: random forest.

We first evaluated our ability to discriminate between genes that were upregulated *vs.* downregulated during aging in neutrophils (**Figure 5A**). Our machine-learning models assigned genes to the correct transcriptional change with age >64% of the time on held-out testing data (64-81% accuracy depending on the algorithm; **Figure 5B**; **Supplementary Table S4A**), significantly above that of a random classification (50%). Such high accuracy suggests that there are clear biological features that differentiate between genes that tend to be up- *vs.* downregulated with aging. To understand which features were most predictive of age-related gene expression remodeling, we examined predictor contribution derived from our RF and GBM models, both of which provide native evaluation of feature importance in trained models (**Figure 5C**; **Supplementary Table S4B**). Consistent with the tight link between chromatin landscape and gene expression (Consortium, 2012), the average ATAC-seq signal at the promoter (*i.e.* describing how “open” the chromatin is at that specific promoter), as well as the fold difference in ATAC-seq signal between young and old neutrophils (*i.e.* age-related changes in chromatin openness at the promoter) were among the top predictors to discriminate age-related changes in gene expression (**Figure 5C**; **Supplementary Table S4B**). Key predictors also included CG and CpG content in the promoter and gene sequences (**Figure 5C**). This observation is consistent with our previous work identifying promoter CpG content as a top predictive feature of age-related gene expression across four male mouse tissues (Benayoun et al., 2019). Cytosines in CpG configurations are the primary targets of DNA methylation in mammalian cells (Rausch et al., 2020). CpG DNA methylation [DNAme] is catalyzed by DNA-methyltransferases (*i.e.* ‘writers’), can be removed through the action of proteins of the TET family (*i.e.* ‘erasers’), and exert biological consequences through recognition by methyl-CpG binding domain proteins (*i.e.* ‘readers’) (Rausch et al., 2020). Consistent with the notion that the predictive power of CpG content in our models is linked to potential defects in the regulation of CpG methylation, our RNA-seq revealed significant transcriptional downregulation with organismal aging of DNA-methyltransferase encoding gene *Dnmt1* (responsible for propagation of methyl-CpG through DNA replication), demethylation-catalyzing TET encoding genes *Tet1* and *Tet2* (which encode ‘erasers’), as well as methyl-CpG binding domain proteins encoding genes *Mbd4* and *Mbd5* (FDR < 5%; **Supplementary Table S1A**). The presence of putative age-related changes in DNAme patterns is consistent with the notion that DNAme patterns can serve as “aging clocks” (Horvath and Raj, 2018).

Key predictors of age-related gene expression changes also included whether a gene was a target of specific regulators: Stat5a, Mtf2, Sirt6, Foxo1, *etc.* (**Figure 5C**; **Supplementary Table S4B**). Although causality cannot be inferred from machine-learning models, regulators predictive of age-related gene expression provide a list of candidate drivers or mediators of gene programs related to organismal age in neutrophils. Only a subset of top TF predictors were themselves differentially regulated with aging according to our RNA-seq (*e.g. Stat5a, Mtf2, Nod2*; **Figure S6G**; **Supplementary Table S1A**). Thus, it is possible that the activity of TF predictors without RNA-level regulation may occur through differential post-translational age-related regulation. Interestingly, Stat5a mediates the effects of GM-CSF on granulocyte differentiation (Feldman et al., 1997), and is crucial for the survival of mature neutrophils (Kimura et al., 2009). Although the role of Mtf2 in neutrophils has not been studied, Mtf2 interacts with Jarid2, another of our top predictors, as part of the PRC2 complex to promote repressive H3K27 methylation (Zhang et al., 2011), mediating the recruitment of PRC2 to unmethylated CpGs specifically (Perino et al., 2018). Nod2 is a pattern recognition receptor that recognizes muramyl dipeptide, the minimal common motif of bacterial peptidoglycans (Girardin et al., 2003). Although Nod2 is not a transcription factor, its activation has been shown to have profound transcriptional effects on gene expression, with target genes including cytokines, chemokines, and regulators of NF-κB signaling (Billmann-Born et al., 2011; Jeong et al., 2014). Indeed, accumulating evidence suggests that Nod2 is an important contributor of neutrophil-mediated innate immunity (Ekman and Cardell, 2010; Jeong et al., 2014). Sirt6 is a histone deacetylase which has been tightly linked to the regulation of mammalian aging longevity (Kanfi et al., 2012; Peshti et al., 2017). Sirt6 can limit inflammation by deacetylating the promoters of NF-kB target genes (Kawahara et al., 2009), and by reducing cytokine production (Lappas, 2012). Consistently, *Sirt6* knockout mice have massive liver inflammation associated to neutrophil infiltration (Xiao et al., 2012). Thus, changes in activity for top predictive TFs may drive aspects of age-related transcriptional remodeling.

Since we identified profound differences across multiple “omics” layers between male and female neutrophils, we next asked whether sex-dimorphic gene expression could be predicted from genomic information, local chromatin states and/or TF regulation (**Figure 6A**). Importantly, as an additional feature, we also included the genomic location of a gene (*i.e.* whether it was located on a sex chromosome or an autosome). Our machine-learning models assigned genes to the correct sex-based expression bias >70% of the time on held-out testing data (70-78% accuracy depending on the algorithm; **Figure 6B**; **Supplementary Table S4C**), consistent with the notion that sex-biases genes share a common regulatory signature. To understand which features were most predictive of male- or female-biased gene expression, we again examined predictor contribution from trained RF and GBM models (**Figure 6C**; **Supplementary Table S4D**). Importantly, being located on a sex chromosome (X or Y) was a poor predictor of sex-dimorphic gene expression (166^th^ combined rank position in predictor importance; **Supplementary Table S4D**), consistent with the fact that direct genetic context is not a crucial driver of sex-dimorphic gene expression (**Figure S1E**). Similar to our age-related models, the average promoter ATAC-seq signal and the fold change between females vs. males in ATAC-seq signal were top also predictors for sex-biased gene expression (**Figure 6C**; **Supplementary Table S4D**). The GC/CpG content of promoter and genes sequences also showed strong predictive power, consistent with the sex-dimorphic expression of DNAme regulators (*i.e.* male-biased expression for *Dnmt1* and *Tet3*; **Supplementary Table S1B**).

Finally, we asked which TFs were most predictive of sex-dimorphic gene expression in neutrophils (**Figure 6C**; **Supplementary Table S4D**). Being a known target of Foxm1, Sirt6, Mtf2 or Mybl2 (also known as b-Myb) was highly predictive of male *vs.* female neutrophil-biased gene expression, suggesting that these regulators may drive aspects of the sex-dimorphic neutrophil biology. Similar to above, only a small subset of top predictors from our sex-dimorphic machine-learning models were themselves significantly regulated at the transcriptional level (e.g. *Foxm1*, *Irf8*; **Figure S6H**; **Supplementary Table S1B**). Although its role hasn’t been specifically explored in neutrophils, Foxm1 is known to regulate the expression of number of cell cycle-related genes (Chen et al., 2013; Grant et al., 2013) and is highly expressed in late committed neutrophil precursors (Kim et al., 2017). The transcription factor b-Myb, is also linked to the regulation of cell cycle-related genes (Zhan et al., 2012), and can modulate differentiation and cell fate decision of myeloid progenitors (Baker et al., 2014). Thus, top predictors of sex-dimorphic neutrophil gene expression Foxm1 and b-Myb might be linked to the observed sex-dimorphic expression of cell cycle and chromatin-related genes (**Figure 3, S4**). Being a target of Sirt6 was also predictive of sex-dimorphic gene expression, which is in line with sex-dimorphic phenotypes in both Sirt6 knock-out and overexpression mice (Kanfi et al., 2012; Peshti et al., 2017). In addition, in line with our lipidomic observation of sex-dimorphism in neutrophil lipid storage (**Figure 3F**), Sirt6 is also tightly linked to lipid metabolism (Jung et al., 2019; Yao et al., 2017), and can limits lipid droplet accumulation in macrophages foam cells (He et al., 2017). In addition, Sirt6 activity can be directly regulated by free fatty acids (*e.g.* myristic, oleic and linoleic acid) (Feldman et al., 2013). Consistently, lipidomics showed that female neutrophils have higher levels of free fatty acids, including FFA(14:0) and FFA(18:2) (*i.e.* myristic and linoleic acid), and may thus have higher basal Sirt6 activity levels (**Supplementary Table S1E,F**). Thus, our machine-learning analysis reveals candidate regulators that may mediate sex-dimorphic phenotypes in neutrophils and will deserve further investigation for their role in neutrophil biology.

### Primary granule-related genes are differentially regulated in females *vs.* males, correlating with differences in basal and elicited release of serum neutrophil elastase

Intriguingly, when analyzing the top genes with the largest RNA-seq fold-difference between male *vs.* female neutrophils, we noticed that a number of genes encoding key components of primary neutrophil granules showed male-biased expression (*e.g. Elane*, *Mpo*, *Prtn3* and *Ctsg* were among the top 25 genes with the largest fold-difference between male and female neutrophils; **Supplementary Table S1B**), suggesting potential differences in granule loading between sexes. The ability of neutrophils to mount an effective defense against pathogens lies in the proteases stored in intracellular granules, so we then used our datasets to investigate potential differences in granule biogenesis (**Figure 7A-B, S7A-D**). Neutrophils store a number of proteases and antimicrobial proteins in 3 main types of granules: (i) primary/azurophilic granules, (ii) secondary/specific granules, and (iii) tertiary/gelatinase granules. To unbiasedly test the hypothesis that neutrophil granule biology is differentially regulated between sexes, we compiled a list of granule components from the literature (**Supplementary Table S5**). Consistent with our first observation, GSEA revealed a robust trend for increased RNA expression of primary (but not secondary or tertiary) granule-related genes in male *vs.* female neutrophils, regardless of age (**Figure 7A-B, S7A-D**).

**Figure 7:**
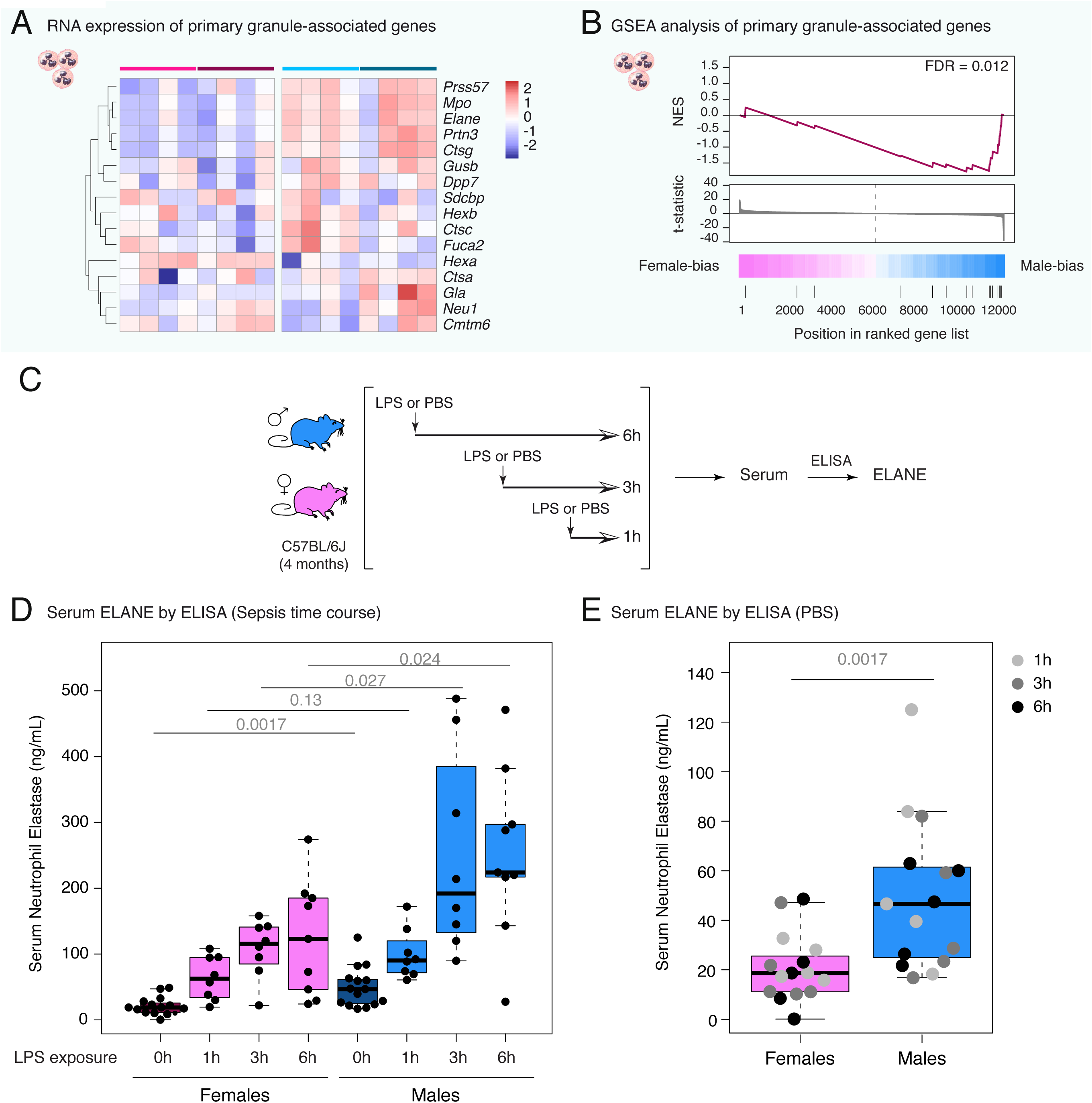
Male neutrophils express higher levels of primary granule genes, correlating with increased serum ELANE levels in control and septic mice. (**A**) Heatmap of normalized gene expression for primary (azurophilic) granule-related gene expression in our RNA-seq dataset. Also see **Supplementary Figure S7**. (**B**) GSEA analysis of primary (azurophilic) granule-related gene expression reveals biased expression to male neutrophils. (**C**) Setup scheme for serum ELANE measurement in control and sepsis-like mice. (**D-E**) Analysis of ELANE protein levels in mouse serum by ELISA. For simplicity’s sake, all PBS-injected animals are reported as “0h of LPS exposure” in (D), and are replotted with time-based color-coding in (E). Significance in non-parametric Wilcoxon rank-sum test.

Neutrophil degranulation is a key aspect of the response of neutrophils to pathogens (Lacy, 2006), and mice without a functional copy of *Elane*, which encodes neutrophil elastase, have increased mortality upon sepsis than controls (Belaaouaj et al., 1998). However, excessive levels of circulating neutrophil elastase are also thought to be deleterious, amplifying septic shock (Barbosa da Cruz et al., 2019) and acute respiratory distress syndrome (Wang et al., 2009). Based on our transcriptomic data, we hypothesized that higher expression of *Elane* in neutrophils could lead to increased circulating levels of neutrophil elastase, both in basal conditions and upon an external immune challenge. To test this hypothesis, we obtained female and male young adult C57BL/6J mice, and injected them with sterile PBS or LPS, a major component of gram-negative bacteria cell wall, for 1-6 hours, to mimic a sepsis-like challenge (**Figure 7C**). Indeed, LPS injection is a simple model to mimic microbial exposure upon septic injury (Raduolovic et al., 2018). Interestingly, males showed significantly higher levels of serum neutrophil elastase at 3 and 6 hours post LPS exposure (**Figure 7D**), as well as in the PBS-injected non-septic controls (**Figure 7D,E**). Thus, males had overall higher serum protein levels of Elane, both in the absence and in the presence of an immune challenge, which is in line with higher *Elane* gene expression in our neutrophil RNA-seq.

Finally, we then took advantage of a published dataset of Data-Independent Acquisition [DIA] proteomics in human neutrophils from 68 healthy donors (Grabowski et al., 2019). Since the biological sex of human donors was not specified, we used the reported protein levels of DDX3Y, a protein encoded on the Y chromosome, as a proxy for the likelihood that the sample was derived from a male donor. Consistent with our mouse neutrophil RNA-seq and our serum ELISA analysis, we found significant positive correlation between levels of DDX3Y (*i.e.* likelihood of sample coming from a male donor) and expression of primary granule proteins ELANE, MPO and CTSG (**Figure S7E-G**), although not for PRTN3. Thus, our results suggest that neutrophil derived from healthy human males may also have higher protein expression levels for key components of primary granules. These observations suggest that, at least for these components, transcriptomic trends are predictive of protein-level trends, and that the male-bias in primary granule components observed in our mouse transcriptomics data may be conserved in humans.

## Discussion

### A resource for the study of neutrophils across sex and lifespan

To understand how aging and biological sex influence neutrophil biology, we have generated transcriptomic, metabolomic, lipidomic and epigenomic datasets using primary bone marrow neutrophils from young and old, female and male animals. To our knowledge, this dataset is the largest multi-omic dataset for the study of neutrophils and we believe it will be a resource to the scientific community. Specifically, it is one of the rare cases to include both sexes and organismal aging, rather than focusing on only one sex or one age group (usually young adults). Thanks to the inclusion of these variables, we observed that (i) neutrophils are extremely sex-dimorphic, with potential consequences on the systemic innate immune response and that (ii) organismal age leads to large scale transcriptomic and epigenomic remodeling of neutrophils. Interestingly, when analyzed separately, males had > 10-fold more significantly regulated genes with age than females, suggesting that the molecular trajectory of neutrophil aging is different between sexes.

Importantly, we already utilized this “multi-omic” resource, and predicted males to have increased release of neutrophil elastase based on neutrophil gene expression patterns. Indeed, we observed that male mice had increased amounts of serum neutrophil elastase both in control and sepsis-like conditions. Excess release of neutrophil elastase is known to exacerbate inflammation and cause tissue damage (Chua and Laurent, 2006). Thus, this resource will help open new avenues of research into neutrophil biology and identify candidate mediators that underlie sex differences in the immune compartment throughout life.

### Chromatin metabolism as a profoundly regulated process in neutrophils across sex and age

Our analyses revealed that transcriptional regulation of chromatin-related pathways is a hallmark of neutrophil aging, and a key aspect with sex-dimorphic regulation. Further, these transcriptional changes were linked to overall differences in chromatin architecture in neutrophils. Far from being a mere regulatory layer for gene expression, neutrophil chromatin organization has profound implications on NETosis, the process by which neutrophil chromatin is extruded in response to immune challenge (Sollberger et al., 2018). Consistent with the unique biological function of chromatin in neutrophils, several studies have reported that neutrophil chromatin organization profoundly diverges from that of other cell types (Chen et al., 2016; Denholtz et al., 2020; Zhu et al., 2017). Our analyses suggest that female neutrophils have an overall “looser” chromatin structure, which is associated to the transcriptional upregulation of a number of transposable elements [TEs]. Conversely (and regardless of sex), neutrophils from old individuals show signs of increased baseline chromatin condensation accompanied by reduced TE transcription. This contrasts with reports of age-related TE derepression across other cell types (Lai et al., 2019), and may reflect the uniqueness of neutrophil biology (*i.e.* overall transcriptional repression, chromatin compaction as a biological function, *etc.*). Thus, it will be important to determine whether these structural changes lead to differences in NETosis between individuals of both sexes and throughout lifespan, with mechanistic implications in aberrant chronic age-related inflammation (Franceschi and Campisi, 2014).

### Machine-learning as a powerful candidate-identification tool

By using machine-learning, we showed that both age-regulation and sex-dimorphism in gene expression can be predicted by an array of features, spanning DNA sequence information, local chromatin accessibility, and putative regulation by candidate TFs. We leveraged information about genes known to have regulatory impact on transcriptional landscapes so as to maximize our likelihood to identify potential drivers of observed age- and sex-regulation patterns. The high accuracy of our trained model supports the notion of coordinated differences between (i) genes that are induced *vs.* repressed in neutrophils during organismal aging, and (ii) genes that are expressed in a sex-dimorphic manner. Among important predictors of transcriptional states, we identified a number of regulators whose activity change during organismal aging may underlie “omic” profile remodeling throughout life, such as Foxo1, Sirt6 or Mtf2. We also identified putative mediators of sex-biased gene expression of neutrophils, such as Foxm1, Mybl2/b-Myb and Irf8. To note, machine-learning does not provide information about causality of relationships. However, important predictors represent top candidate mediators that could help shape neutrophil molecular or functional landscapes with aging or as a function of sex. Thus, it will be important to elucidate the potential role of these predictors on neutrophil biology in order to understand the emergence of age-related and sex-dimorphic phenotypes.

### Neutrophils as a mediator cell type for sex-differences in immune aging

Accumulating evidence has shown that, even outside of the reproductive organs, mammalian biology is extremely sex-dimorphic (Sampathkumar et al., 2020). This is especially relevant to organismal aging, with large sex-differences in baseline lifespan and in response to interventions that successfully extend the health- and lifespan in laboratory mice (Austad and Bartke, 2015). Although this has not been broadly investigated with aging, pioneering studies have started to compare the molecular landscapes of male and female cells, revealing profound sex-dimorphism in gene regulation across cells and tissues, including immune cells (Gal-Oz et al., 2019; Marquez et al., 2020; Qu et al., 2015; Yen and Kellis, 2015). Consistently, accumulating evidence has shown that immune responses differ between males and females (Angele et al., 2014; Buskiewicz et al., 2016; Kovats, 2015).

Sex-dimorphism in immune outcomes is exemplified in the current health crisis, with men representing the majority of severe cases and deaths from COVID-19 (Jin et al., 2020; Petrilli et al., 2020; Scully et al., 2020; Serge et al., 2020). Interestingly, NET release is thought to drive aspects of inflammatory disease, sometimes leading to tissue damage (Papayannopoulos, 2018; Sollberger et al., 2018), and might contribute to severe COVID-19 disease (Barnes et al., 2020; Zuo et al., 2020). Thus, molecular differences between male and female neutrophils are likely to contribute to an array of pathologies and could constitute targets to optimize immune responses throughout life and help tailor therapeutic strategies to men and women.

## Materials and methods

### Mouse husbandry

All animals were treated and housed in accordance to the Guide for Care and Use of Laboratory Animals. All experimental procedures were approved by the University of Southern California’s Institutional Animal Care and Use Committee (IACUC) and are in accordance with institutional and national guidelines. For ‘omics’ analyses, male and female C57BL/6Nia mice (4 and 20 months old animals) were obtained from the National Institute on Aging (NIA) colony at Charles Rivers. For the neutrophil purity and sepsis assays in young animals, male and female C57BL/6J mice (3-4 months old animals) were obtained from Jackson Laboratories. Both Charles Rivers and Jackson Laboratories have specific-pathogen-free (SPF) facilities. Animals were acclimated at the SPF animal facility at USC for 2-4 weeks before any processing. All animals were euthanized between 9-11am for ‘omics’ experiments and flow cytometry. For the sepsis model experiments, animals were injected in the morning, and euthanized between 4-6pm. In all cases, animals were euthanized using a “snaking order” across all groups to minimize batch-processing confounds (as well as potential counfounds due to neutrophil cellular “age”). All animals were euthanized by CO2 asphyxiation followed by cervical dislocation.

### Isolation of primary neutrophils from mouse bone marrow

The long bones of each mouse were harvested and kept on ice in D-PBS (Corning) supplemented with 1% Penicillin/Streptomycin (Corning) until further processing. Muscle tissue was removed from the bones, and the bone marrow from cleaned bones was collected into clean tubes (Amend et al., 2016). Red blood cells from the marrow were removed using Red Blood Cell Lysis buffer (Miltenyi Biotech #130-094-183), according to the manufacturer’s instructions, albeit with no vortexing step to avoid unscheduled neutrophil activation. Neutrophils were isolated from other bone marrow cells using magnetic-assisted cell sorting (Miltenyi Biotech kit #130-097-658). Viability and yield were assessed using trypan blue exclusion and an automated COUNTESS cell counter (Thermo-Fisher Scientific). Purified cells were pelleted at 300g and snap-frozen in liquid nitrogen until processing for RNA, lipid or metabolite isolation.

### Neutrophils purity estimate by flow cytometry

We used a cohort of 5 young female and 5 young male C57BL/6J mice to estimate potential differences in neutrophil purity with respect to sex. After an Fc-blocking step (Miltenyi Biotech #130-092-575), MACS-purified neutrophils were then stained using APC-Ly6G (Invitrogen #17-9668-80) and Vioblue-Cd11b (Miltenyi Biotech #130-113-238) at a 1:50 dilution according to the manufacturer’s instructions. Stained cells were then analyzed by flow cytometry on a MACS Quant Analyzer 10 (Miltenyi Biotech). Flow cytometry results were processed using the FlowLogic V7 software. Purity of MACS-isolated bone marrow neutrophils in young adult male and female animals are reported in **Figure S2A-C**.

### Estimate of neutrophil proportion in bone marrow in young female and male mice

We used a cohort of 5 young female and 5 young male C57BL/6J mice to estimate potential differences in neutrophil proportion within nucleated bone marrow cells with respect to sex. An aliquot of bone marrow cell suspension was obtained after bone marrow extraction and red blood cell lysis (see above). Cell composition analysis was obtained using the Hemavet 950FS at the USC Leonard Davis School of Gerontology mouse phenotyping core. Percentage of cells was used instead of absolute cell numbers to avoid confounding results due to animal size.

### RNA purification and RNA-seq library preparation

For RNA isolation, frozen cell pellets were resuspended in 1mL of Trizol reagent (Thermo-Fisher), and total RNA was purified following the manufacturer’s instructions. RNA quality was assessed using the Agilent Bioanalyzer platform at the USC Genome Core using the RNA Integrity Number (RIN). 500ng of total RNA was subjected to ribosomal-RNA depletion using the NEBNext rRNA Depletion Kit (New England Biolabs), according to the manufacturer’s protocol. Strand specific RNA-seq libraries were then constructed using the SMARTer Stranded RNA-seq Kit (Clontech), according to the manufacturer’s protocol. Libraries were quality controlled on the Agilent Bioanalyzer 2100 platform at the USC Genome Core, before multiplexing the libraries for sequencing. Paired-end 75bp reads were generated on the Illumina NextSeq500 platform at the SC^2^ Core at CHLA.

### RNA-seq analysis pipeline

Paired-end 75bp reads were processed using Trimgalore 0.4.4 (github.com/FelixKrueger/TrimGalore) (i) to retain only high-quality bases with phred score > 15, and (ii) to eliminate biases due to priming by hard clipping the first 6 bases of each read. Only pairs with both reads retaining a length of > 45bp after trimming were retained for further processing. Trimmed reads were mapped to the mm10 genome reference using STAR 2.5.0a (Dobin et al., 2013). Read counts were assigned to genes from the UCSC mm10 reference using subread 1.5.3 (Liao et al., 2014) and were imported into R to perform differential gene expression analysis. Based on general RNA-seq processing guidelines, only genes with mapped reads in at least 6/16 RNA-seq libraries were considered to be expressed and retained for downstream analysis. Due to high sample-to-sample variability, we used surrogate variable analysis to estimate and correct for unwanted experimental noise (Leek and Storey, 2007). R package ‘sva’ v3.34 was used to estimate surrogate variables, and the removeBatchEffect function from ‘limma’ v3.42.2 was used to regress out the effects of surrogate variables and RNA-integrity differences (RIN scores) from raw read counts. The ‘DESeq2’ R package (DESeq2 1.26.0) was used for further processing of the RNA-seq data in R (Love et al., 2014). Genes with a false discovery rate < 5% were considered statistically significant and are reported in **Supplementary Table S1A-B**.

### Dimensionality reduction for exploratory data analysis

To perform Multidimensional Scaling (MDS) analysis, we used a distance metric between samples based on the Spearman’s rank correlation value (1-Rho), which was then provided to the core R command ‘cmdscale’. Dimensionality reduction was applied to pre-normalized data, as described in relevant sections.

### Putative Protein-Protein Interaction [PPI] network analysis

Genes with significant regulation according to our DEseq2 analysis (FDR < 5%) were used as input for network analysis with NetworkAnalyst 3.0 (Zhou et al., 2019). For age-related gene regulation, significant genes and DEseq2 calculated log_2_(fold-change) were used as input for a single network analysis. For sex-dimorphic gene regulation, the female- and male-biased gene lists were used as separate inputs for the analysis. To avoid network clusters due to sex-chromosome encoded genes, only significant autosomal genes were included in the sex-dimorphism gene expression networks. In both cases, we had NetworkAnalyst use PPI information derived from IMEx/InnateDB data, a knowledgebase specifically geared for analyses of innate immune networks (Breuer et al., 2013), to construct putative PPI networks. In each case, the largest subnetwork determined by NetworkAnalyst was used in figures and analyses.

### Functional enrichment analysis (transcriptomics)

We used the Gene Set Enrichment Analysis (GSEA) paradigm (Subramanian et al., 2005) through the ‘phenotest R package (1.34.0). Gene Ontology term annotation were obtained from ENSEMBL through Biomart (Ensembl 99; downloaded on 2020-04-10), and gene-term association with only author statement support (GO evidence codes ‘TAS’ and ‘NAS’) or unclear evidence (GO evidence code ‘ND’) were filtered out. Other annotations were obtained from the Molecular Signature Database v7.0 (*e.g.* Reactome, KEGG) (Liberzon et al., 2011; Subramanian et al., 2005) and the Harmonizome Database (*e.g.* ChEA, JASPAR, ENCODE, GEO TF targets; accessed 2020-03-25) (Rouillard et al., 2016). To improve target coverage, we summarized putative transcription factor [TF] targets, including FOXO TF targets compiled in our previous study (Benayoun et al., 2019), so as to have a reduce unique and non-redundant list of TF targets summarized from all these sources. The DEseq2 t-statistic was used to generate the ranked list of genes for functional enrichment analysis, both for the sex and aging effects. For ease of reading, only the top 10 most significant pathways with negative NES and top 10 most significant pathways with positive NES are shown in figures if more than that pass the FDR < 5% significance threshold. All significant gene sets are reported in **Supplementary Tables S2** (aging) and **S3** (sex). In parallel, functional enrichment was also independently assessed using the Ingenuity Pathway Analysis portal, using genes with a significant male or female bias (FDR < 5%).

### Chemicals

LC-MS-grade solvents and mobile phase modifiers were obtained from Fisher Scientific (water, acetonitrile, methanol, methyl tert-butyl ether) and Sigma−Aldrich (ammonium acetate).

### Metabolite and lipid extraction from neutrophils

Metabolites and lipids were extracted from neutrophil cell pellets and analyzed in a randomized order. Extraction was performed using a biphasic separation protocol with ice-cold methanol, methyl tert-butyl ether (MTBE) and water (Contrepois et al., 2018). Briefly, 300μL of methanol spiked-in with 54 deuterated internal standards provided with the Lipidyzer platform (SCIEX, cat #5040156, LPISTDKIT-101) was added to the cell pellet, samples were vigorously vortexed for 20 seconds and sonicated in a water bath 3 times for 30 seconds on ice. Lipids were solubilized by adding 1000μL of MTBE and incubated under agitation for 1h at 4°C. After addition of 250μL of ice-cold water, the samples were vortexed for 1 min and centrifuged at 14,000g for 5 min at 20°C. The upper phase containing the lipids was then collected and dried down under nitrogen. The dry lipid extracts were reconstituted with 300μL of 10 mM ammonium acetate in 9:1 methanol:toluene for analysis. The lower phase containing metabolites was subjected to further protein precipitation by adding 4 times of ice-cold 1:1:1 isopropanol:acetonitrile:water spiked in with 17 labeled internal standards and incubating for 2 hours at −20°C. The supernatant was dried down to completion under nitrogen and re-suspended in 100μL of 1:1 MeOH:Water for analysis.

### Untargeted LC-MS metabolomics

#### Data acquisition

Metabolic extracts were analyzed four times using hydrophilic liquid chromatography (HILIC) and reverse phase liquid chromatography (RPLC) separation in both positive and negative ionization modes as previously described (Contrepois et al., 2015). Data were acquired on a Thermo Q Exactive plus mass spectrometer equipped with a HESI-II probe and operated in full MS scan mode. MS/MS data were acquired on pool samples consisting of an equimolar mixture of all the samples in the study. HILIC experiments were performed using a ZIC-HILIC column 2.1×100 mm, 3.5μm, 200Å (Merck Millipore) and mobile phase solvents consisting of 10mM ammonium acetate in 50/50 acetonitrile/water (A) and 10 mM ammonium acetate in 95/5 acetonitrile/water (B). RPLC experiments were performed using a Zorbax SBaq column 2.1 × 50 mm, 1.7 μm, 100Å (Agilent Technologies) and mobile phase solvents consisting of 0.06% acetic acid in water (A) and 0.06% acetic acid in methanol (B). Data quality was ensured by (i) injecting 6 and 12-pool samples to equilibrate the LC-MS system prior to run the sequence for RPLC and HILIC, respectively, (ii) sample randomization for metabolite extraction and data acquisition, and (iii) checking mass accuracy, retention time and peak shape of internal standards in every samples.

#### Data processing

Data from each mode were independently analyzed using Progenesis QI software v2.3 (Nonlinear Dynamics). Metabolic features from blanks and that didn’t show sufficient linearity upon dilution were discarded. Only metabolic features present in >33% of the samples in each group were kept for further analysis and missing values were imputed by drawing from a random distribution of small values in the corresponding sample (Tyanova et al., 2016).

#### Metabolic feature annotation

Annotation confidence levels for each metabolite were provided following the Metabolomics Standards Initiative (MSI) confidence scheme. Peak annotation was first performed by matching experimental m/z, retention time and MS/MS spectra to an in-house library of analytical-grade standards (Level 1). Remaining peaks were identified by matching experimental m/z and fragmentation spectra to publicly available databases including HMDB (http://www.hmdb.ca/), MoNA (http://mona.fiehnlab.ucdavis.edu/) and MassBank (http://www.massbank.jp/) using the R package ‘MetID’ (v0.2.0) (Shen et al., 2019) (Level 2). Briefly, metabolic feature tables from Progenesis QI were matched to fragmentation spectra with a m/z and a retention time window of ±15 ppm and ±30 s (HILIC) and ± 20 s (RPLC), respectively. When multiple MS/MS spectra match a single metabolic feature, all matched MS/MS spectra were used for the identification. Next, MS1 and MS2 pairs were searched against public databases and a similarity score was calculated using the forward dot–product algorithm which takes into account both fragments and intensities. Metabolites were reported if the similarity score was above 0.4. Level 3 corresponds to unknown metabolites.

### Targeted lipidomics with the Lipidyzer platform

#### Data acquisition

Lipid extracts were analyzed using the Lipidyzer platform that comprises a 5500 QTRAP System equipped with a SelexION differential mobility spectrometry (DMS) interface (SCIEX) and a high flow LC-30AD solvent delivery unit (Shimazdu). A full description of the method is available elsewhere (Contrepois et al., 2018). Briefly, the lipid molecular species were identified and quantified using multiple reaction monitoring (MRM) and positive/negative switching. Two acquisition methods were employed covering 10 lipid classes; method 1 had SelexION voltages turned on, while method 2 had SelexION voltages turned off. Lipidyzer data were reported by the Lipidomics Workflow Manager (LWM) software which calculate concentration in nmol/g for each detected lipid as average intensity of the analyte MRM/average intensity of the most structurally similar internal standard MRM multiplied by its concentration. Data quality was ensured by i) tuning the DMS compensation voltages using a set of lipid standards (SCIEX, cat #5040141) after each cleaning, more than 24 hours of idling or 3 days of consecutive use, ii) performing a quick system suitability test (QSST) (SCIEX, cat #50407) before each batch to ensure acceptable limit of detection for each lipid class, iii) sample randomization for lipid extraction and data acquisition, and iv) triplicate injection of lipids extracted from a reference plasma sample (SCIEX, cat #4386703) at the beginning of the batch.

#### Data pre-processing

Lipids detected in less than 66% of the samples in each group were discarded and missing values were imputed in each class by drawing from a random distribution of small values in the corresponding sample (Tyanova et al., 2016).

### Differential analysis of metabolomics and lipidomics data

Metabolomics and lipidomics datasets were first normalized to the total protein content as determined by BCA assay (Pierce #23225) to account for differential starting material quantity. Then, Variance Stabilizing Normalization was applied to the data using ‘limma’ 3.42.2, as recommended by previous studies (Jauhiainen et al., 2014; Li et al., 2016). Differential analysis for metabolomic or lipidomic features was performed using ‘limma’ in R. Features with a false discovery rate (FDR) < 5% were considered statistically significant.

For analysis of the lipidomics data by lipid class (**Supplementary Figure S1D**; **Supplementary Table S1F**), lipids were summarized by class after BCA and VSN corrections. To determine regulation at the class level, because of the small number of analyzed classes, a basic linear model approach was used through base R ‘lm’ function, and FDR correction was applied using the base R ‘p.adjust’ function. Lipid classes with a false discovery rate (FDR) < 5% were considered statistically significant.

### Functional enrichment analysis for metabolomics

Since only 1 metabolite that was significantly regulated with aging, we focused on analyzing differentially regulated metabolite sets with respect to sex. To analyze the directional regulation of metabolic pathways from untargeted metabolomics, we used R package ‘MetaboAnalystR’ 2.0.4 (Chong and Xia, 2018) to perform Phenotype Set Enrichment analysis (PSEA) (Ried et al., 2012) of KEGG metabolic pathways. Using the ‘mummichog’ method (Li et al., 2013), we provided an input table of metabolic peaks represented by mass over charge ratios (m/z) and retention time, limma-derived p-value and t-scores, and analysis mode (negative or positive ion) to have maximum sensitivity for the functional enrichment analysis of untargeted metabolomic data. All significant gene sets (FDR < 5%) are reported in **Supplementary Table S3F**.

### Integrated functional enrichment analysis for RNA-seq and metabolomics using IMPaLA

Based on the differential analyses results in RNA-seq and metabolomics, we focused on analyzing differentially regulated genes and metabolites with respect to sex. To provide an integrated view of RNA-seq and metabolomics results, we took advantage of the IMPaLA paradigm (Kamburov et al., 2011). We used the IMPaLA v12 web interface (http://impala.molgen.mpg.de/), which evaluates overrepresentation of genes and metabolites across 5055 annotated pathways from 12 databases, with default parameters. For the joint analysis, we used genes with FDR < 5% with respect to biological sex, using gene symbols as input IDs. For metabolic features, we selected only the manually validated species (Levels 1 and 2) with FDR < 5% with respect to biological sex. Finally, we also selected lipid species FDR < 5% with respect to biological sex. Importantly, we used HMDB IDs as input IDs for the metabolomic/lipidomic side of the analysis, thus analyzing only features with correponding HMDB IDs. Joint analysis of significant features with a male bias, or, separately, those with female bias, were uploaded separately to the server (server access on 06-30-2020). All pathways with an overall combined FDR < 5% are reported in **Supplementary Table S3H**.

### Functional enrichment analysis for lipidomics

Since there were no significant difference in lipid composition with aging, we focused on analyzing differentially regulated lipid sets with respect to sex. The Lipid Ontology (LION) website was used to perform functional enrichment analysis of lipids (Molenaar et al., 2019). Lipid features with FDR < 5% with a male or female bias were uploaded to the server (access on 03-24-2020). For ease of reading, only the 10 most significant pathways with male and with female bias are shown in **Figure 3F**. All significant LION terms are reported in **Supplementary Table S3G**.

### Neutrophil ATAC-seq library generation

An independent cohort of C57BL/6Nia mice was used to assess age- and sex-related differences in chromatin architecture using ATAC-seq. To assay potential differences in the chromatin landscape of mouse bone marrow neutrophils across age and sex, we used the omni-ATAC protocol starting from 50,000 MACS-purified cells (Corces et al., 2017). Libraries were quality controlled on the Agilent Bioanalyzer 2100 platform at the USC Genome Core before pooling. Libraries were multiplexed and sequenced on the Illumina HiSeq-Xten platform as paired-end 150bp reads at the Novogene Corporation (USA).

### ATAC-seq preprocessing pipeline

Paired-end ATAC-seq reads were adapter-trimmed using NGmerge0.2 (Gaspar, 2018), which clips overhanging reads in a sequence-independent fashion and has been recommended for use in ATAC-seq preprocessing. Trimmed paired-end were mapped to the mm10 genome build using bowtie2 (2.2.6) (Langmead and Salzberg, 2012). After alignment, PCR duplicates were removed using the ‘rmdup’ function of samtools (1.5). To minimize analytical artifacts from uneven sequencing depth between biological samples/libraries, libraries were randomly downsampled to the same depth for downstream analyses using PicardTools (2.20.3) or samtools (1.5). We used the peak finding function of HOMER (4.11) to identify ATAC-seq accessible regions (Heinz et al., 2010).

### Differential accessibility analysis with ATAC-seq

We extracted a normalized read count matrix from our the downsampled bam files over merged HOMER regions using R package ‘Diffbind’ 2.14 (Ross-Innes et al., 2012). We used this extracted matrix for downstream analyses. Because of sample-to-sample variability, we used surrogate variable analysis (Leek and Storey, 2007) to estimate and correct for experimental noise, similar to the RNA-seq analysis. R package ‘sva’ 3.34 was used to estimate surrogate variables, and the ‘removeBatchEffect’ function from ‘limma’ 3.42.2 was used to regress out the effects of surrogate variables, PCR duplicate content and variation in reads mapping to peaks according to Diffbind (Fraction of reads in peaks). The ‘DESeq2’ R package (1.26.0) was used for further processing of ATAC-seq data. Regions with a false discovery rate < 5% were considered statistically significant.

### Chromatin architecture analysis from ATAC-seq datasets

To analyze the underlying chromatin architecture deifferences between neutrophils isolated from different biological groups, we used the NucleoATAC paradigm (Schep et al., 2015). Since NucleoATAC requires high sequencing depth to reliably measure nucleosome profiles, we pooled depth-matched reads from each biological group to attain this depth as recommended by the authors of the software (Schep et al., 2015). We extracted similar metrics from the NucleoATAC output to what was described in previous studies using this software to investigate chromatin architecture (King et al., 2018).

### Repetitive element transcription analysis

To evaluate potential differential regulation of transposable elements, we used the ‘analyzeRepeats.pl’ functionality of the HOMER software to count STAR-mapped reads over repetitive elements (Heinz et al., 2010). To allow for whole library normalization, reads mapping over transcript were also counted using the same HOMER script. Read counts were imported into R to estimate differential repeat expression using the ‘DESeq2’ (1.26.0) R package (Love et al., 2014).

### Feature extraction for machine-learning analysis

For each significantly age-regulated or sex-dimorphic gene in neutrophils according to DEseq2 (FDR < 5%), we extracted a number of features associated to that gene. First, we took advantage of our ATAC-seq data to evaluate the chromatin architecture of neutrophils (see above). For each gene, we extracted (i) the average promoter accessibility by ATAC-seq in FPKM across all 4 conditions (promoters were defined as −500bp;+150bp with regards to annotated transcriptional start sites in mm10 build according to HOMER), as well as (ii) the average log_2_(Fold Change) in accessibility between old and young neutrophils (age-regulation models) **<u>or</u>** the average log_2_(Fold Change) in accessibility between female and male neutrophils (sex-dimorphism models). Second, we took advantage of gene sets coverage known of predicted targets of transcriptions factors: ChEA, ENCODE, JASPAR and GEO TF perturbation information from the Harmonizome database (Rouillard et al., 2016) and transcriptional targets of FOXO transcription factors from GEO experiments compiled in our previous study (Benayoun et al., 2019). To reduce extraneous features, we engineered a TF target feature following these steps: (i) TF targets were summarized from all putative sources, (ii) only TFs with evidence of expression in primary neutrophils according to RNA-seq were retained as features, and (ii) only TFs with at least 25 targets across significant age-related genes (age-regulation models) or sex-dimorphic genes (sex-dimorphism models) were retained. This process yielded 357 (age-regulation models) and 249 (sex-dimorphism models) TF target sets used as features for model training. Finally, we included several DNA sequence features to each gene: the percentage of CG nucleotide and the percentage of CpG dinucleotide in promoters and exons, computed using HOMER. To note, HOMER was only able to provide information for 3,421 of the 3,653 genes with significant age regulation, and 1,636 of the 1,734 genes with significant sex-dimorphic regulation. Finally, to determine whether location of the gene on autosomes vs. chromosomes was a key predictive factor for sex-dimorphic gene expression, we also included for the sex-dimorphic gene expression models a feature encoding whether the gene is located on autosomes or sex chromosomes (X or Y).

### Machine-learning analysis for age-regulated or sex-dimorphic genes

We trained machine-learning classification models for 2 questions: (i) models for age-regulated gene expression, and (ii) models for sex-dimorphic gene expression. For (i), age-regulation machine-learning models were built to learn whether up- or down-regulated genes with could be discriminated using genomic features and predict potential master regulators. For (ii), sex-dimorphism machine-learning models were built to learn whether female and male biased genes could be discriminated using genomic features and predict potential master regulators.

In both cases, we built classification models using 7 classification algorithms as implemented through R package ‘caret’ (caret 6.0-86). Auxiliary R packages were used with ‘caret’ to implement neural networks (NNET; ‘nnet’ 7.3-13), random forests (RF; ‘randomForest’ 4.6-14), gradient boosting (GBM; ‘gbm’ 2.1.5), radial kernel support vector machines (SVM; kernlab 0.9-29), sparse Linear Discriminant Analysis (LDA; ‘sparseLDA’ 0.1-9), conditional inference Trees (cTree; ‘party’ 1.3-4), Regularized Logistic Regression (LogReg; ‘LiblineaR’ 2.10-8). ‘Caret’ was allowed to optimize final model parameters on the training data using 10-fold cross validation. Accuracies, sensitivities, specificities and AUC (using package ‘pROC’ 1.16.2) for all trained classifiers were estimated using a test set of randomly held out 1/3 of the data (not used in the training phase) obtained using the ‘createDataPartition’ function. Feature importance estimation was only done using tree-based RF and GBM methods, since other algorithms do not natively allow for it. The feature importance was computed and scaled by ‘caret’ for the RF and GBM models.

### Reanalysis of DIA proteomics data from human neutrophils

We obtained DIA proteomics results from the supplemental material of a recent study (Grabowski et al., 2019). We used normalized expression values from the article’s supplemental material and used the reported “donor-specific Spectronaut protein expression values” for further analysis. To exclude potential confounds linked to disease, we only used data from the 68 healthy donors [HD] and excluded the data from diseased patients. Since the article did not report the biological sex of donors, we used detected expression levels of DDX3Y, a Y-chromosome encoded protein, as a proxy for the likelihood of the sample belonging to a male donor. We then examined rank correlation statistics between protein expression of DDX3Y compared to that of human orthologs of top male-biased primary granules genes from our mouse RNA-seq data (*i.e.* ELANE, MPO, PRTN3 and CTSG). Significance of the Spearman rank correlation is reported.

### Mouse sepsis model through intra-peritoneal LPS injection

Young adult (3-4 months) C57BL/6J mice were exposed to LPS, a pathogen-associated molecular pattern (PAMP) found in the cell-wall of Gram-negative bacteria, to elicit a sepsis-like response (Raduolovic et al., 2018). Briefly, mice were injected intra-peritoneally with 2µg of LPS per gram of body weight, using a sterile LPS stock (Sigma #L5293) diluted in PBS, and control animals were injected with sterile PBS (Raduolovic et al., 2018). Animals were monitored hourly for up to 6 hours after injection for the occurrence and severity of endotoxemia (Raduolovic et al., 2018). After euthanasia by CO_2_ asphyxiation and cervical dislocation at 1, 3 and 6 hours post injection, blood was collected from the heart. To obtain serum for downstream analysis, the blood was left to clot for ≥1 hour at room temperature in MiniCollect Serum SeparatorTubes (Greiner Bio-One). The serum was then separated from the clot using centrifugation at 2,000g for 10 minutes, and frozen at −80°C until further use.

### Quantification of serum neutrophil elastase (ELANE) by ELISA

Quantitative evaluation of circulating ELANE levels was performed from serum. ELISA quantification of serum ELANE levels was performed using Abcam’s Mouse Neutrophil Elastase SimpleStep ELISA Kit (ab252356) according to the manufacturer’s instructions. Technical replicates from the same sample were averaged as one value before statistical analysis and plotting.

### Data Availability

Sequencing data has been submitted to the Sequence Read Archive (SRA) accessible through BioProject PRJNA630663. Untargeted metabolomics data was uploaded to metabolomics workbench DataTrack ID 2089. The lipidomics data is available as **Supplemental Table S6**.

### Code Availability

The analytical code will be made available on the Benayoun lab Github repository (https://github.com/BenayounLaboratory/). All R code was run using R version 3.6.0-3.6.3.

## Acknowledgements

We would like to thank Dr. Daniel Campo and Suchi Patel at the USC genome core for assistance in NGS-library quality control on the Agilent Bioanalyzer platform; Dr. Fan Li at the SC2 Core at CHLA for help with sequencing of transcriptomic libraries on the Illumina NextSeq550. We thank Dr. Todd Morgan and Gerald Navarette from the USC Leonard Davis School of Gerontology mouse phenotyping core for assistance with CBC analyses on the Hemavet 950FS. We acknowledge the use of the HPC resource at USC for computational analyses. We thank Dr. Changhan Lee (USC), Dr. Kelvin Yen (USC) and Dr. Helen Goodridge (Cedars-Sinai Medical Center) for helpful insights and feedback on our study. Finally, we thank members of the Benayoun and Vermulst labs for helpful discussions and feedback.

## Author contributions

S.T. and B.A.B. designed the study. R.L., S.T., N.K. and B.A.B. isolated and/or processed neutrophil samples for “omic” profiling. K.C. experimentally processed samples for metabolomic and lipidomic profiling and pre-processed the resulting data. K.C. and M.E. performed the manual validation/identification of significant metabolites from the untargeted metabolomics data. R.L. and B.A.B. conducted the “sepsis-like” experiment. B.A.B. performed data analyses. B.A.B wrote the manuscript with input from all the authors. All authors edited and commented on the manuscript.

## Supplementary material

### Legends to Supplementary Figures

**Figure S1:**
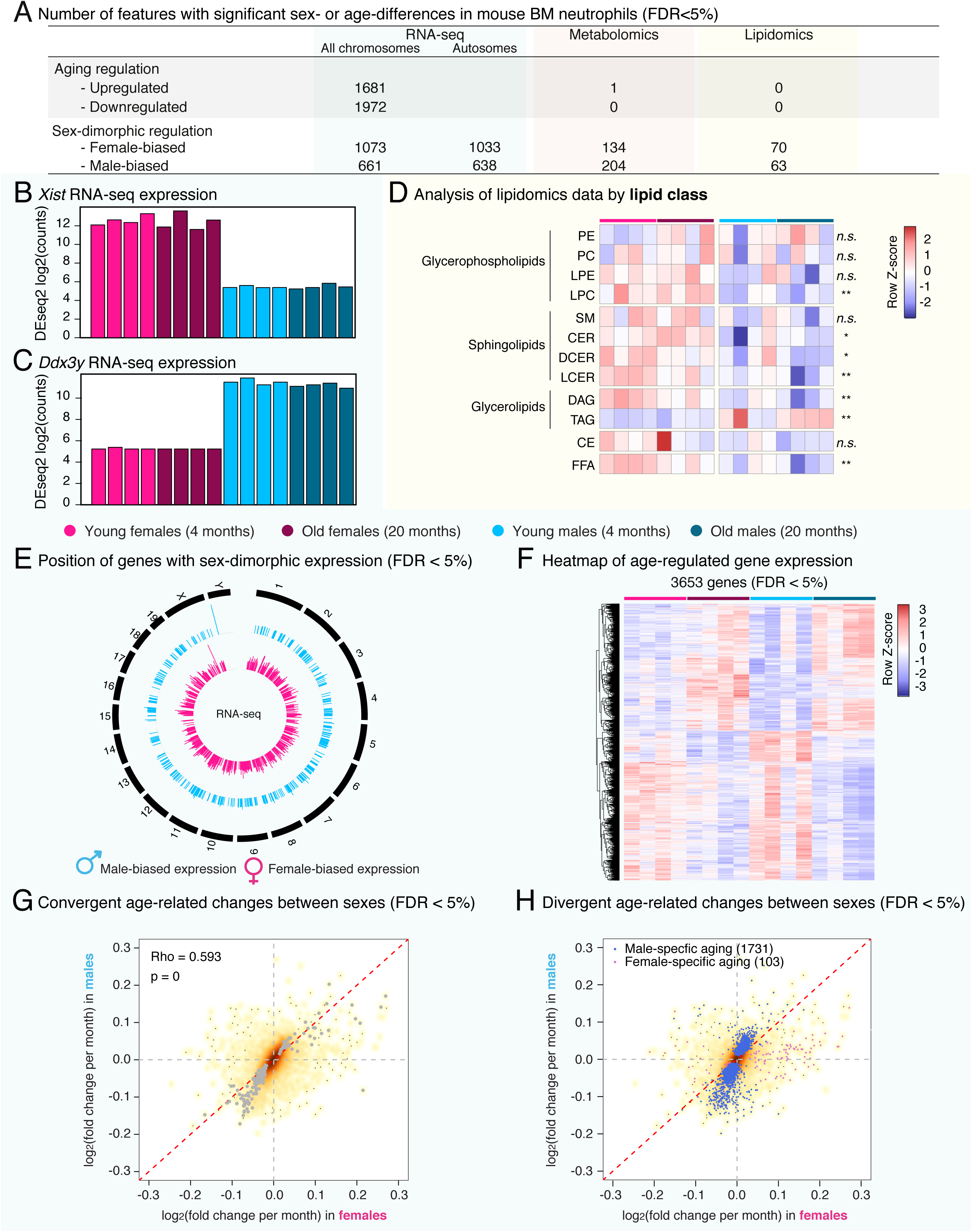
A multi-omic analysis of primary mouse bone marrow neutrophils during aging and with respect to sex (continued). (**A**) Table of significant “omic” features as a function of organismal age and sex in our datasets. For the analysis of sex-dimorphism in gene expression, the number of significant genes located on autosomes (*i.e.* not on chromosomes X or Y) is also reported. (**B-C**) Barplot of DESeq2-normalized log2 counts for *Xist* (B) and *Ddx3y* (C), showing the expected pattern between male and female samples. (**D**) Heatmap of lipidomic changes summarized by lipid classes. Significance of difference as a function of age or sex was evaluated using a linear model approach, and the significance of the sex coefficient is reported on the line of the heatmap. * : p < 0.05; ** : p < 0.01; n.s. : p ≥ 0.05. See also Supplementary **Table S1F**. (**E**) Circular genome plot of the positions of genes with significant sex-biased in neutrophil gene expression (FDR < 5%). (**F**) Heatmap of significant age-regulated genes (DESeq2 FDR < 5%). (**G-H**) Correlation plot of age-related gene expression change in female *vs.* male neutrophils from RNA-seq, showing genes with (G) significant and concordant age-regulation in both sexes at FDR 5%, or (H) genes with divergent age-regulation between sexes (FDR < 5% in one sex, and FDR >15% in the other). Spearman Rank correlation (Rho), and significance of this correlation are reported in (G).

**Figure S2:**
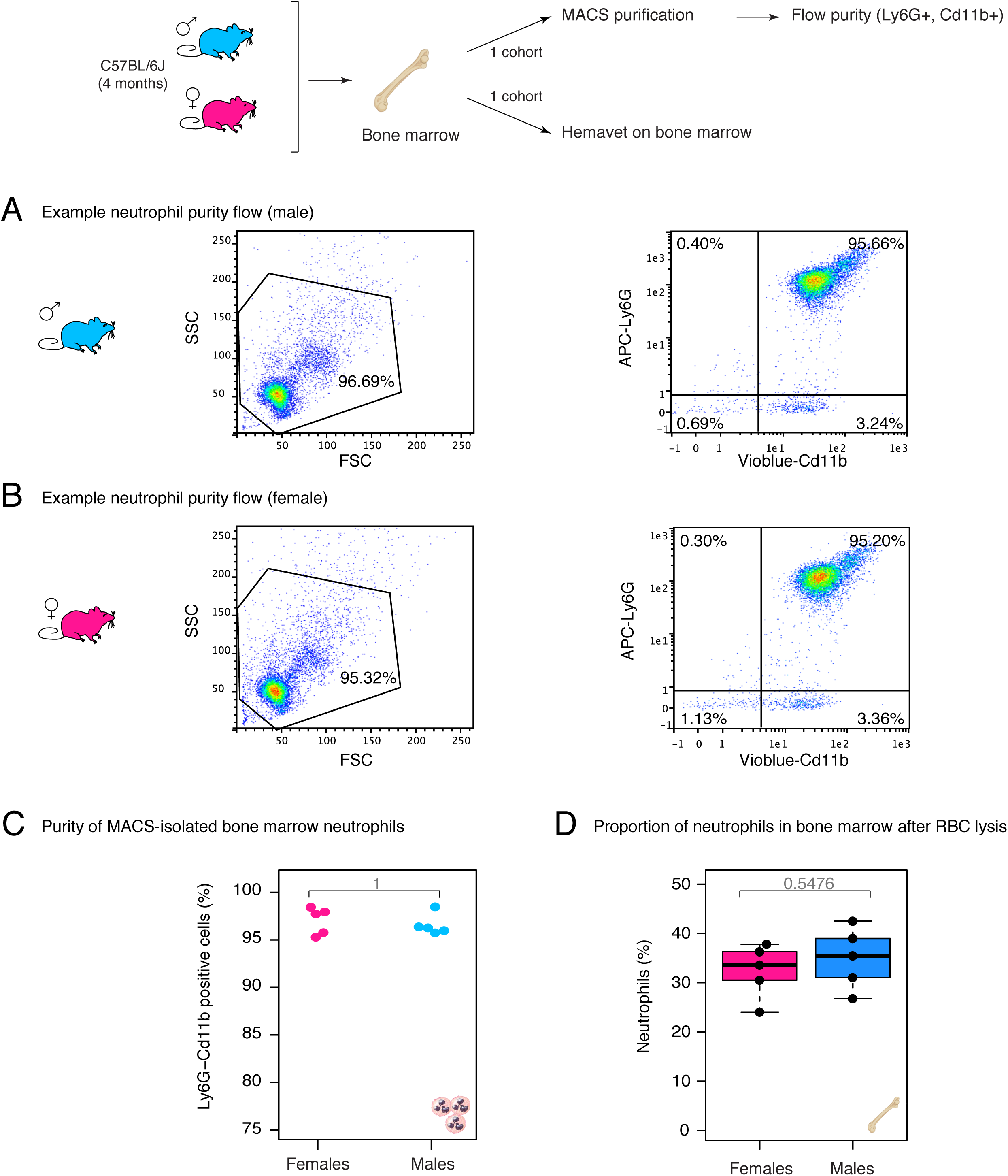
Bone marrow neutrophil purity is not impacted by animal sex. (**A-B**) Representative flow cytometry analysis of bone-marrow neutrophils purified using MACS from young male (A) or female (B) mice. Neutrophils are expected to be double positive for Cd11b and Ly6G. (**C**) Neutrophil purity from a test cohort of male vs. female mice (n = 5), determined as the Cd11b+ Ly6G+ population. (**D**) Proportion of neutrophils among bone marrow nucleated cells according to the Hemavet 950FS. P-value for (C) and (D) derived from non-parametric Wilcoxon rank-sum test.

**Figure S3:**
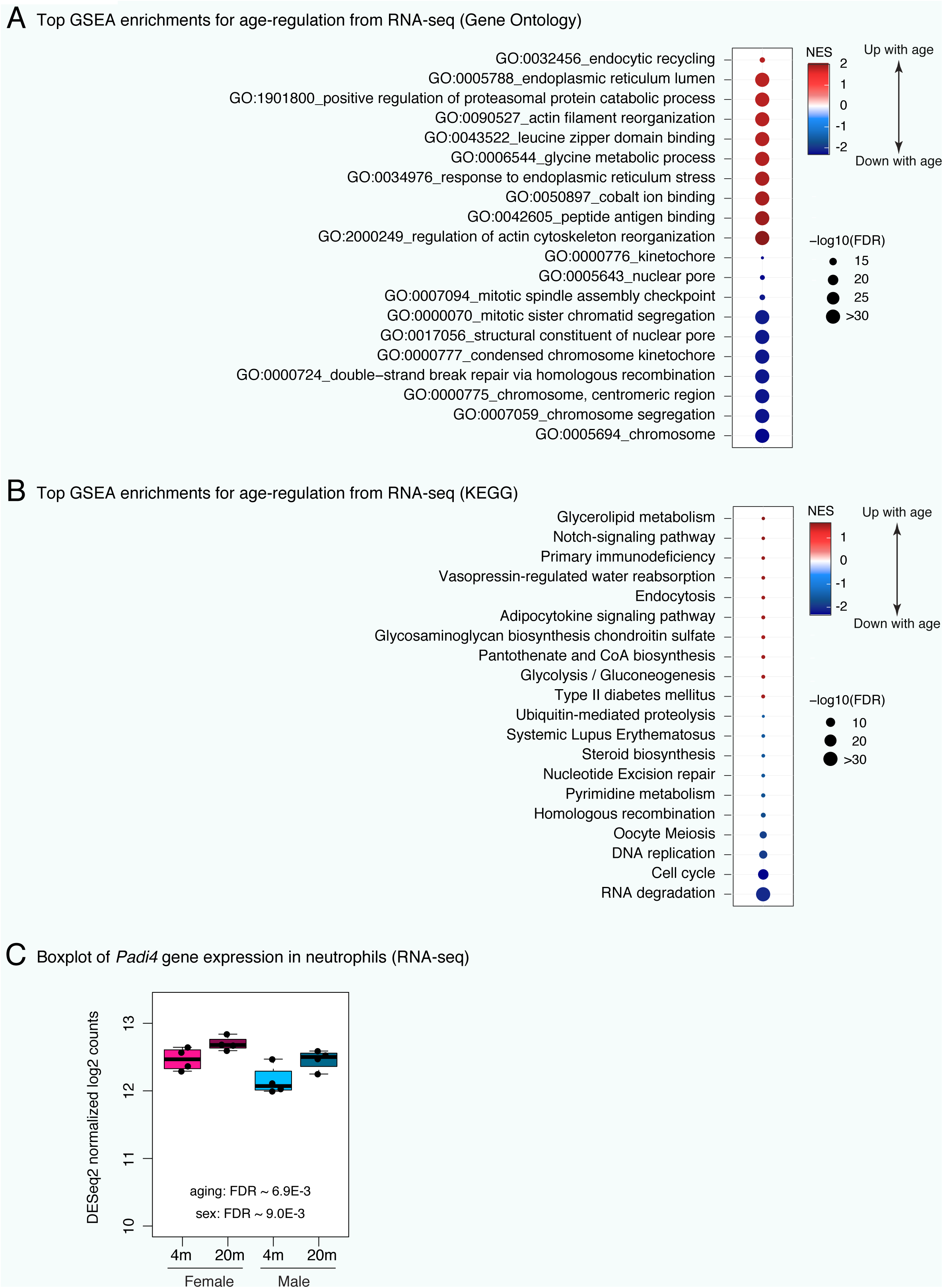
Misregulated pathways in bone marrow neutrophils during aging reveals downregulation of chromatin-related pathways (continued). (**A-B**) Top enriched gene sets from Gene Ontology (A) and KEGG (B) using GSEA for differential RNA expression. Only the top 10 most up- and top 10 most downregulated gene sets are plotted for readability. Full lists and statistics available in **Supplementary Table S2.** Also see **Supplementary Figure S3**. All shown pathways and genes such as FDR < 5%. NES: Normalized Enrichment Score (for GSEA analysis). FDR: False Discovery Rate. (**C**) Boxplot of *Padi4* transcriptional levels from RNA-seq. Significance: DESeq2 FDR values for regulation as a function of aging or biological sex.

**Figure S4:**
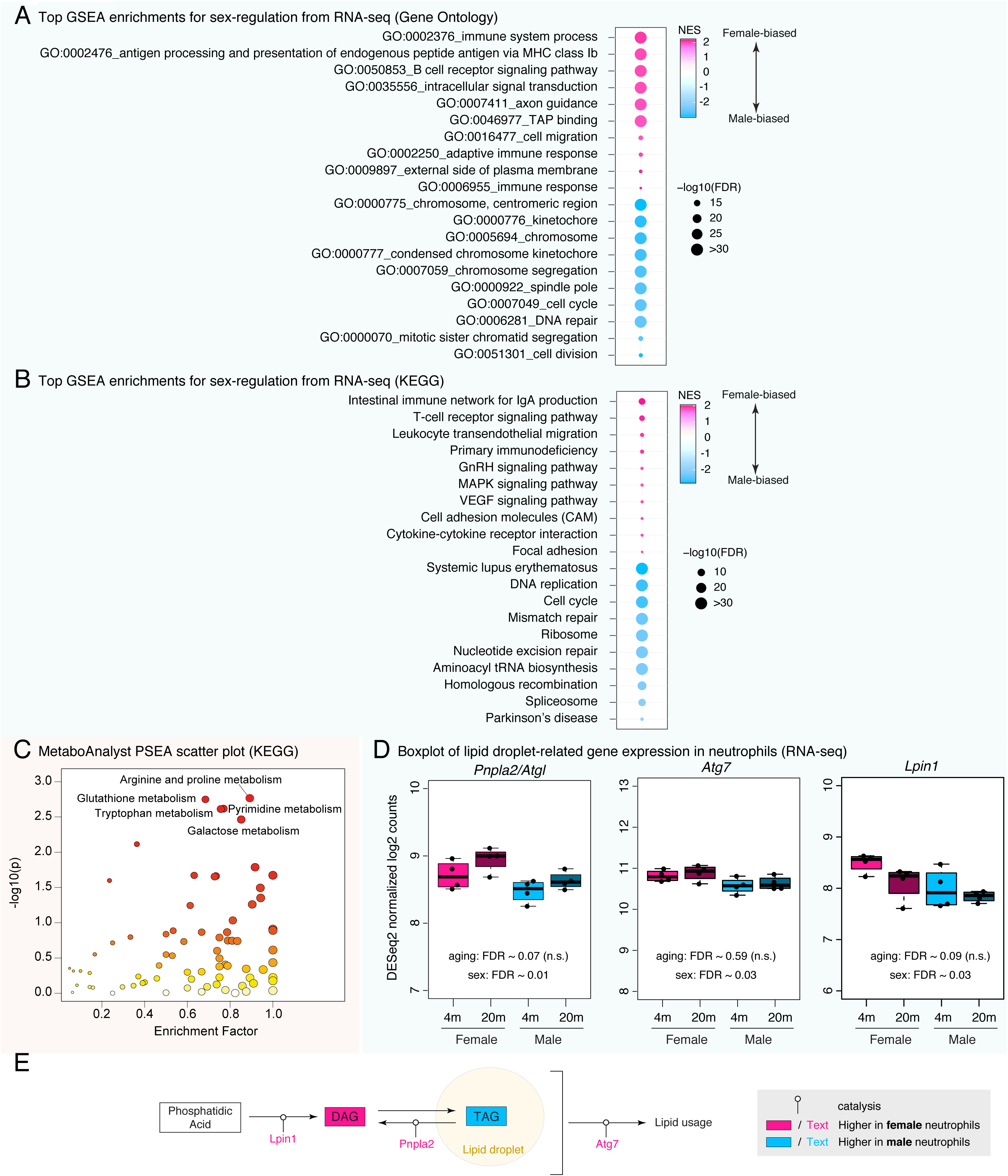
Sex-dimorphic pathways in bone marrow neutrophils reveals differential regulation of chromatin-related pathways (continued). (**A-B**) Top enriched gene sets from Gene Ontology (A) and KEGG (B) using GSEA for differential RNA expression as a function of sex. Only the top 10 most up- and top 10 most downregulated gene sets are plotted for readability. Full lists and statistics available in **Supplementary Table S2.** Also see **Supplementary Figure S3**. All shown pathways and genes such as FDR < 5%. (**C**) MetaboAnalyst PSEA scatterplot for KEGG pathways from metabolomics. NES: Normalized Enrichment Score (for GSEA analysis). FDR: False Discovery Rate. (**D**) Boxplot of transcriptional levels from our RNA-seq for *Pnpla2/Atgl*, *Atg7* and *Lpin1*, which are sex-dimorphic lipid metabolism-related genes. Significance: DESeq2 FDR values for regulation as a function of aging or biological sex. (**E**) Summary scheme of discussed lipid usage in female vs. male neutrophils based on metabolomic and transcriptomic data. Catalyzed reactions are derived from Wikipathway WP3901 (Lipid droplet metabolism) and (Riffelmacher et al., 2017).

**Figure S5:**
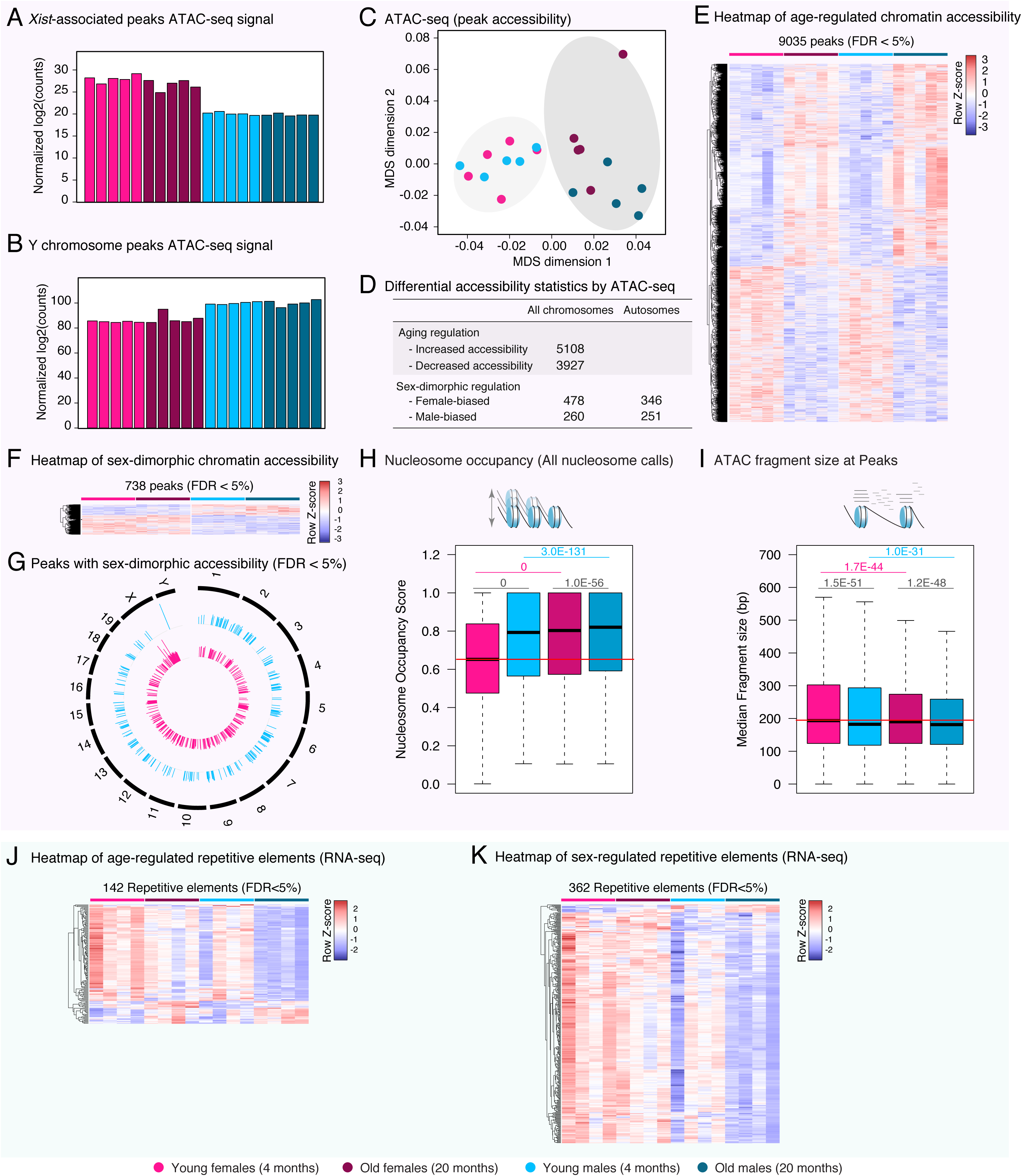
ATAC-seq analysis reveals age- and sex-related differences in the chromatin architecture of bone marrow neutrophils (continued). (**A-B**) Barplot of DESeq2-normalized log2 counts at ATAC-seq peaks associated to *Xist* (A) or situated on the Y chromosome (B), showing the expected pattern between male and female samples. (**C**) Multidimensional Scaling analysis results of chromatin accessibility from ATAC-seq. (**D**) Table of significantly differentially accessible ATAC-seq peaks as a function of organismal age and sex. The number of significant peaks located on autosomes (*i.e.* not on chromosomes X or Y) is also reported. (**E-F**) Heatmap of accessibility at peaks with significant age-regulated (E) or sex-dimorphic (F) change in accessibility (FDR < 5%). (**G**) Circular genome plot of the genomic positions of peaks with significant sex-biased in neutrophil ATAC-seq (FDR < 5%). (**H**) Boxplot of nucleosome occupancy as calculated by NucleoATAC for all called nucleosomes across groups. Also see **Figure 4E**. (**I**) Boxplot of median ATAC-seq fragment length at ATAC-seq peaks across experimental groups. The horizontal red line in panels (**H,I**) shows the median value in neutrophils from young females for ease of comparison. Significance in non-parametric Wilcoxon rank-sum tests are reported for panels (**H,I**). Black p-values represent differences between male *vs.* female neutrophils in each age group; pink (blue) p-values represent age-related differences in female (male) neutrophils. (**J-K**) Heatmap of repetitive elements with significant age-regulated (J) or sex-dimorphic (K) change in transcription from RNA-seq (FDR < 5%).

**Figure S6:**
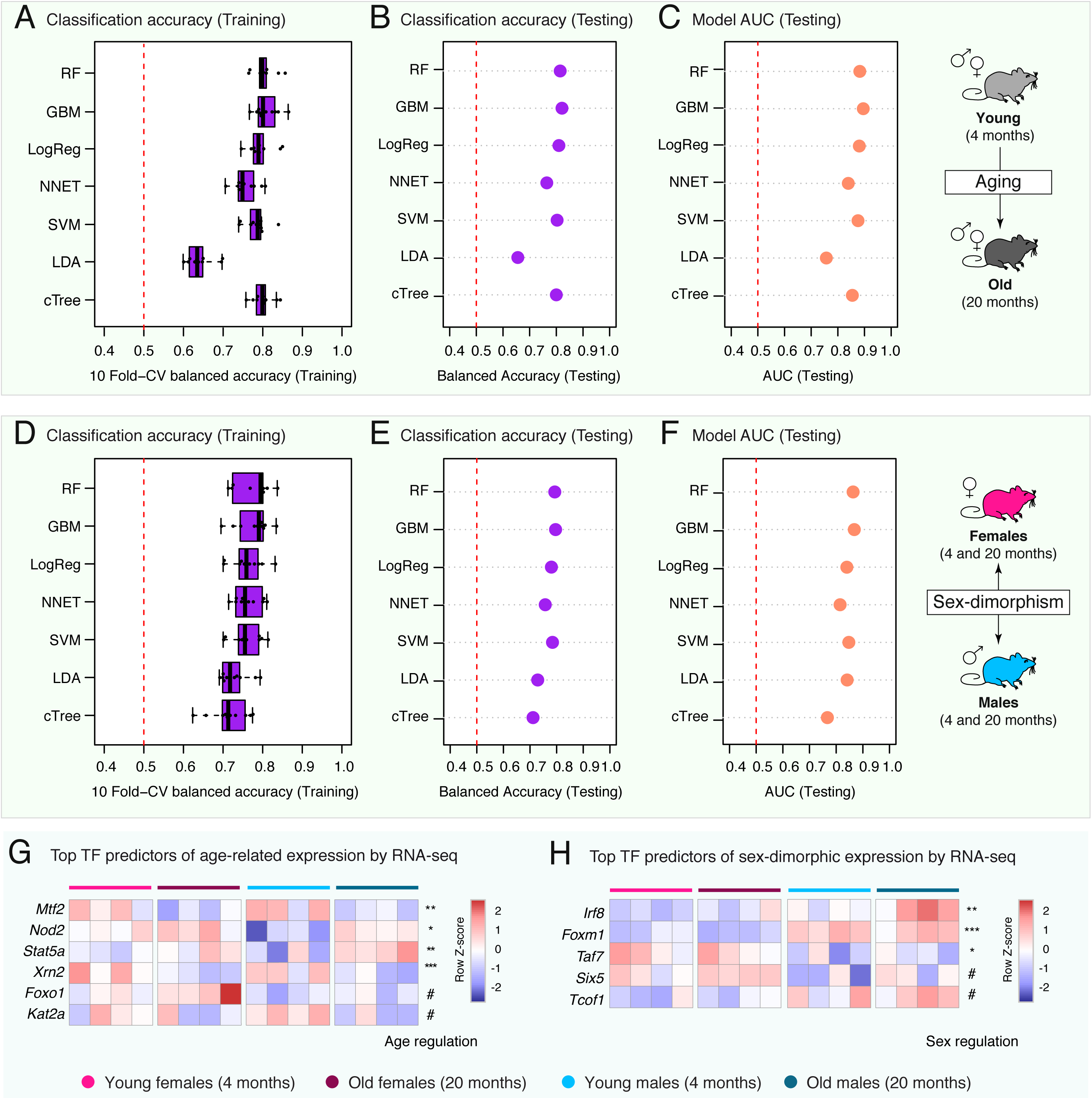
Machine-learning analysis reveals that age-regulated and sex-dimorphic gene expression can be predicted by genomic and epigenomic features. (**A-C**) Age-related machine-learning model performance metrics: balanced classification accuracy over the 10 cross-validation folds during model training (A), balanced classification accuracy on held-out testing data (B), and Area Under the Curve [AUC] on held-out testing data (C). Also see **Supplementary Table S4A**. (**D-F**) Sex-dimorphism machine-learning model performance metrics: balanced classification accuracy over the 10 cross-validation folds during model training (D), balanced classification accuracy on held-out testing data (E), and Area Under the Curve [AUC] on held-out testing data (F). Also see **Supplementary Table S4C**. (**G-H**) Heatmap of RNA-seq expression level for top TF predictors in age-regulated (G) or sex-dimorphic (H) gene expression models with FDR < 10%. DESeq2 FDR is reported on each line, with # : FDR < 0.1; * : FDR < 0.05; ** : FDR < 0.01; *** : FDR < 0.001. FDR: False Discovery Rate.

**Figure S7:**
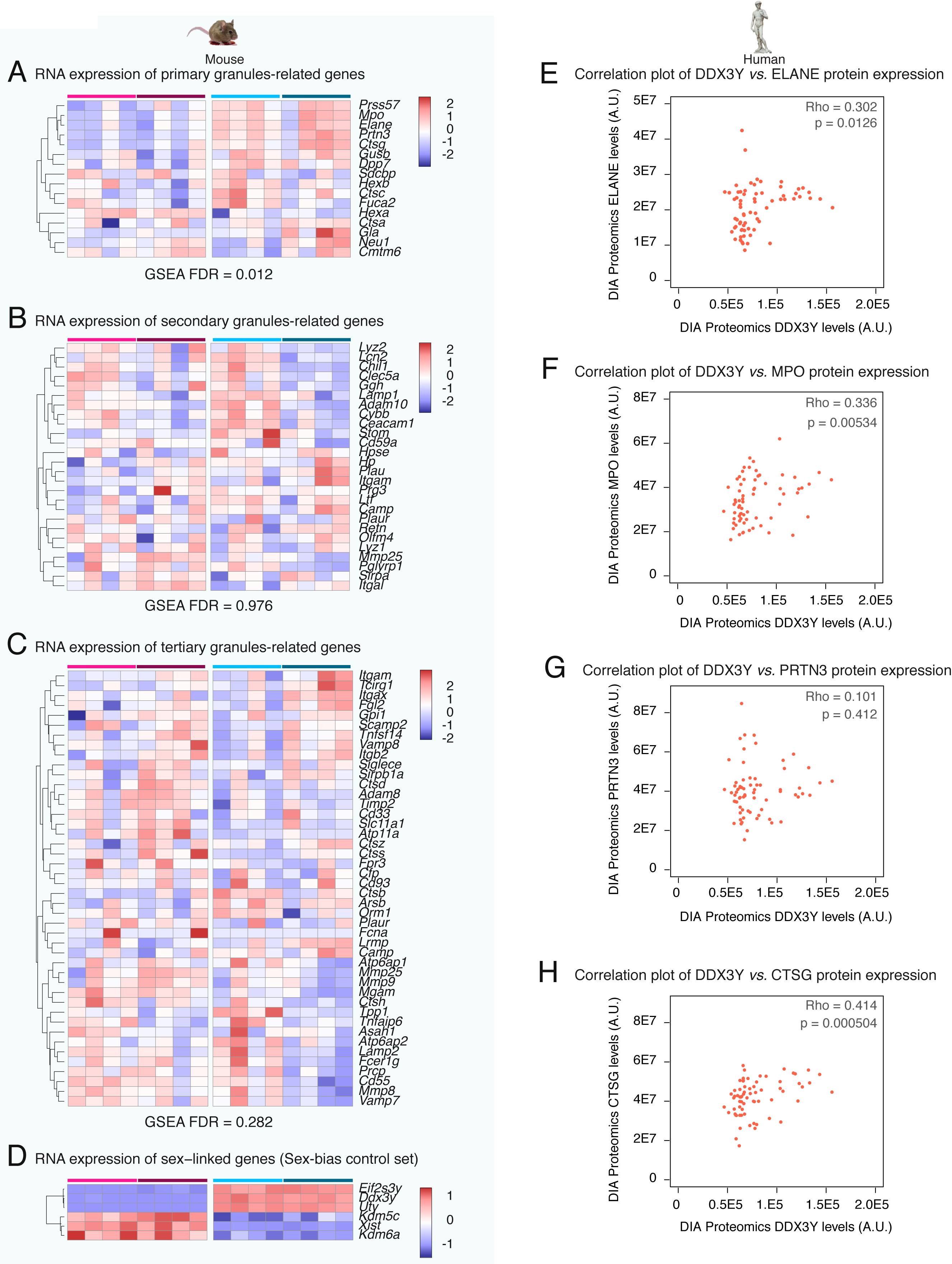
Male neutrophils express higher levels of primary granule genes but not secondary and tertiary granule genes. (**A-C**) Heatmap of normalized gene expression for primary (A), secondary (B) and tertiary (C) granule-related gene expression in our RNA-seq dataset. The estimated False Discovery Rate [FDR] using GSEA for each gene set is reported. (**D**) Heatmap of normalized gene expression for X-linked (*Kdm5c*, *Kdm6a*, *Xist*) and Y-linked (*Uty*, *Ddx3y*, *Eif2s3y*) genes in our RNA-seq dataset. Panel (A) is identical to **Figure 7A** to help direct comparison between sex-bias expression patterns of different granule types as well as canonical sex-biased genes. (**E-H**) Scatterplots of protein levels of DDX3Y (as a proxy for likely of a sample being derived from a male donor), compared to protein levels for primary granule-related proteins ELANE (E), MPO (F), PRTN3 (G) and CTSG (H) in neutrophils from healthy <u>human</u> donors. Normalized data was obtained from the supplementary material of (Grabowski et al., 2019). Estimates of Spearman Rank correlation (Rho) and significance of correlation are reported for each correlation on the scatterplot.

### Supplementary Table Inventory

**Table S1: Table of significantly regulated “omic” features as a function of age or sex in neutrophils.**

(**A**) DEseq2 significantly regulated genes with aging in neutrophils by RNA-seq (FDR <5%). (**B**) DEseq2 significantly regulated genes between male and female neutrophils by RNA-seq (FDR <5%). (**C**) Limma significantly regulated metabolites with aging by untargeted metabolomics (FDR <5%). (**D**) Limma significantly regulated metabolites between male and female neutrophils by untargeted metabolomics (FDR <5%). (**E**) Limma significantly regulated lipids between male and female neutrophils by lipidomics (FDR <5%) (**F**) Linear model analysis by lipid class for regulation in neutrophils by sex or age by lipidomics. (**G**) DEseq2 significantly regulated repetitive elements with aging in neutrophils by RNA-seq (FDR <5%). (**H**) DEseq2 significantly regulated repetitive elements between male and female neutrophils by RNA-seq (FDR <5%).

**Table S2: Table of significantly regulated gene sets with aging in neutrophils by GSEA or IPA.**

(**A**) GSEA MSigDB KEGG (FDR < 5%) for differential regulation with respect to age (RNA-seq). (**B**) GSEA MSigDB Reactome (FDR < 5%) for differential regulation with respect to age (RNA-seq). (**C**) GSEA Gene Ontology (FDR < 5%) for differential regulation with respect to age (RNA-seq). (**D**) GSEA targets of neutrophil-expressed TFs from GEO, ChEA, ENCODE and JASPAR (FDR < 5%) for differential regulation with respect to age (RNA-seq). (**E**) IPA canonical pathway differentially regulated with respect to age (RNA-seq).

**Table S3: Table of significantly regulated gene/feature sets as a function of sex in neutrophils by GSEA, IPA, MetaboAnalyst PSEA or LION.**

(**A**) GSEA MSigDB KEGG (FDR < 5%) for differential regulation with respect to sex (RNA-seq). (**B**) GSEA MSigDB Reactome (FDR < 5%) for differential regulation with respect to sex (RNA-seq). (**C**) GSEA Gene Ontology (FDR < 5%) for differential regulation with respect to sex (RNA-seq). (**D**) GSEA targets of neutrophil-expressed TFs fromGEO, ChEA, ENCODE and JASPAR (FDR < 5%) for differential regulation with respect to sex (RNA-seq). (**E**) IPA canonical pathway differentially regulated with respect to sex (RNA-seq). (**F**) Metaboanalyst PSEA KEGG analysis for differential regulation with respect to sex (metabolomics). (**G**) Integrated analysis of significant sex-biased pathways (FDR < 5%) by IMPaLA (RNA-seq, lipidomics and metabolomics). (**H**) LION functional enrichment analysis for differential regulation with respect to sex (lipidomics).

**Table S4: Table of machine-learning related metrics.**

(**A**) Machine-learning model for aging gene expression (up *vs.* down-regulated with age) performance metrics on held-out testing data. (**B**) Variable importance information for RF and GBM models for aging gene expression (up *vs.* down-regulated with age). (**C**) Machine-learning model for sex-dimorphic gene expression (male *vs.* female bias) performance metrics on held-out testing data. (**D**) Variable importance information for RF and GBM models for sex-dimorphic gene expression (male *vs.* female bias).

**Table S5: Curated lists of proteins found in neutrophil granules.**

(**A**) Curated list of genes encoding proteins found in primary/azurophilic granules according to the literature. (**B**) Curated list of genes encoding proteins found in secondary/specific granules according to the literature. (**C**) Curated list of genes encoding proteins found in tertiary/gelatinase granules according to the literature.

**Table S6: Targeted lipidomics data on SCIEX platform.**

(A) Output of the SCIEX platform for quantification. (B) BCA readings for normalization.

